# Mechanistic prediction of fecal microbiota transplant engraftment and *Clostridioides difficile* colonization resistance with μODE

**DOI:** 10.64898/2026.10.09.758137

**Authors:** Fabio Cumbo, Jayadev Joshi, Bryan Raubenolt, Natalie Katzenmeyer, Jan Claesen, Daniel Blankenberg

## Abstract

Shotgun metagenomics routinely resolves microbial community composition down to the strain level via metagenome-assembled genomes (MAGs), but translating these static catalogs into mechanistic, time-resolved understandings of ecosystem dynamics remains a critical computational bottleneck.

Here, we present μODE, a fully automated framework that converts MAGs into dynamical simulations of community metabolism equipped with a composable, literature-grounded ecology layer, fundamentally designed to serve as a mechanistic hypothesis-testing engine. It employs dynamic flux balance analysis (dFBA) coupled to a shared extracellular pool and integrates a composable ecology layer capturing pH inhibition, time-varying antibiotic pharmacokinetics, bile-acid transformations, and spore dormancy.

To evaluate the core engine, we cross-validated the dFBA machinery to machine precision against an independent reference implementation. We provide native export to the established COMETS engine for third-party verification. Benchmarking against two independent experimentally validated datasets confirmed that while monoculture growth is robustly captured, overflow secretion and directed community composition remain strictly bounded by the flux-routing limitations of max-biomass FBA. However, when applied to rate-modifying ecological feedback, μODE mechanistically and quantitatively reproduces bile-acid-mediated colonization resistance against *Clostridioides difficile*. With a single rate parameter calibrated so that baseline deoxycholate matched its measured physiological concentration, the framework reproduced the uncalibrated antibiotic-induced collapse of deoxycholate, with the calibrated inhibitor concentration coinciding with the pathogen’s independently set inhibitory threshold and driving a ∼44x shift in the simulated pathogen-susceptibility ratio; bile-acid transformation is represented in the composable ecology layer rather than in the genome-scale stoichiometry. Finally, applying μODE end-to-end to a real longitudinal fecal microbiota transplant (FMT) yielding 139 *de novo* genome-scale metabolic models, the framework successfully recovered the clinical direction of treatment. By transparently diagnosing the structural competitive-exclusion bias that limits precise compositional prediction, μODE motivates the necessity of community-objective extensions while transitioning microbiome modeling from descriptive taxonomy toward testable, quantitatively-validated dynamic hypothesis testing, laying the essential mechanistic groundwork for the predictive engineering of safe, targeted ecological therapeutics. μODE is freely available at https://github.com/cumbof/muODE.

---

Over the last two decades, the study of microbial communities has been transformed by sequencing. Amplicon surveys first made it routine to ask *who is there* ^1–4^. Shotgun metagenomics then deepened the answer from marker-gene taxonomy to gene- and path-level functional potential ^5–9^, and metagenome-assembled genomes (MAGs) now reconstruct hundreds of near-complete, often novel, genomes directly from a single complex sample, frequently at strain resolution and without cultivation ^10–13^. A modern gut, soil, ocean, or bioreactor metagenome routinely yields a catalogue of the genomic components of a community of unprecedented completeness ^14–17^.

However, the questions that matter biologically and clinically are not compositional but functional and dynamic: which metabolites does the community actually exchange; which members depend on which others; how is that interaction network reconfigured when the diet changes ^18–20^, when an antibiotic is given ^21–23^, or when a probiotic or a whole donor community is introduced ^24–27^? Answering these requires moving along a specific progression – *who is there* → *what they can do* (functional potential) → *what they are doing* (realized flux in context) → *how it changes over time* (dynamics) – and it is the last two steps, the move from potential to realized, time-resolved function, where automation is weakest.

Currently, genome-scale metabolic models (GEMs) and flux balance analysis (FBA) provide the standard route from a genome to its metabolic capabilities ^28–31^. Community-level extensions of FBA typically compute a single optimal community state (i.e., a steady-state snapshot of who grows on what). While these snapshots are valuable, a great deal of community biology is intrinsically dynamic and cannot be read off a single optimum. For instance, sequential resource use and ecological succession dictate that early colonizers create the specific biochemical niches and cross-fed metabolites that later members exploit ^32–34^. Furthermore, snapshots that assume a fixed medium cannot represent a community that continuously alters its own environment, such as when gut fermenters produce short-chain fatty acids that accumulate, acidify the lumen, and ultimately inhibit the producers themselves ^35–39^. Crucially, the most profound consequences of clinical interventions are often temporal, emergent cascades. The true impact of an antibiotic extends beyond killing susceptible organisms; it can have secondary ecological implications through disrupting cross-feeding partners that suddenly lose an essential exchanged metabolite ^40,41^. Similarly, a fecal microbiota transplant (FMT) is a timed ecological event whose success relies on transient metabolic competition against an established incumbent community ^42–45^. Capturing these emergent behaviors requires dynamic flux balance analysis (dFBA) ^46,47^, a time-stepping scheme that couples each organism’s intracellular optimization to the ordinary differential equations (ODEs) modeling the shared extracellular environment.

Here, we present μODE, a fully automated, reproducible framework that directly converts massive MAG catalogs into predictive, time-resolved community simulations. By employing a native integration engine with saturable Michaelis-Menten kinetics and a shared proteome budget ^48–52^, μODE ensures that phenomena like overflow metabolism, nutrient depletion, and cross-feeding emerge naturally from competition for a shared extracellular pool rather than relying on prescribed, hard-coded rules. Furthermore, we surround this metabolic core with a composable ecology layer, capturing complex environmental interactions such as pH toxicity, bile-acid transformations, spore dormancy, and time-varying antibiotic pharmacokinetics, as well as a perturbation engine to model discrete timed interventions like FMTs and specific gene knockouts ^53–56^.

To rigorously evaluate the framework, we first cross-validate the dynamic engine to machine precision and benchmark it against two independent, controlled *in vitro* datasets to quantitatively delimit its predictive envelope. We then transition to a clinical phenotype, demonstrating its mechanistic capacity to quantitatively reproduce measured *in vivo* bile-acid-mediated colonization resistance against *Clostridioides difficile*. Finally, we apply μODE end-to-end to a longitudinal clinical FMT to diagnose the metabolic drivers of pathobiont suppression. In doing so, we transparently map the structural biases of max-biomass FBA that currently limit exact compositional forecasting. Rather than obscuring these limitations, μODE uses them to redefine the modeling objective: by explicitly separating a composable, literature-grounded ecology layer from the metabolic core, the framework shifts the computational focus away from pure compositional prediction and toward time-resolved, mechanistic hypothesis testing.

## RESULTS

While modern shotgun metagenomics routinely resolves the compositional structure of microbiomes into hundreds of metagenome-assembled genomes (MAGs), translating this static parts list into a mechanistic, time-resolved understanding of community function remains a fundamental bottleneck. Existing constraint-based tools rely predominantly on steady-state snapshots ^31,57^, requiring substantial manual curation and struggling to capture the temporal, environmental feedbacks that govern real ecosystems. To bridge the gap between static functional potential and realized ecosystem dynamics, we developed μODE, a fully automated, end-to-end computational framework for simulating microbial communities.

Unlike existing approaches that rely on prescribed interactions or global steady-state optima, μODE employs a dynamic flux balance analysis (dFBA) engine rooted in the static-optimization approach ^58,59^. Crucially, community members within μODE do not interact directly. Instead, their individual genome-scale metabolic models (GEMs) are coupled solely through a shared, dynamic pool of extracellular metabolites. By integrating saturable Michaelis-Menten uptake kinetics with this compartmentalized architecture, complex ecological phenomena, such as sequential resource depletion, cross-feeding dependencies, and competitive exclusion, are never hard-coded. Rather, they emerge natively from fundamental thermodynamic and mass-balance constraints.

To operate at the scale of modern metagenomics, μODE fully automates the pipeline from raw MAG ingestion to dynamic simulation. The framework executes kingdom-aware reconstruction, structural quality gating, and gap-filling without manual intervention. μODE natively supports two distinct reconstruction backends: a top-down approach utilizing CarveMe (BiGG namespace) ^60,61^ and a bottom-up approach utilizing gapseq (ModelSEED namespace) ^62–64^. This flexibility is critical; as we demonstrate in downstream analyses, gapseq is explicitly required for capturing key anaerobic secretion networks in gut consortia, as the BiGG reference universe currently lacks a connected butyrate efflux pathway.

Finally, because pure metabolism is insufficient to capture the realities of host-associated or environmental microbiomes, we wrapped the metabolic core in a composable ecology layer. This layer introduces critical non-metabolic constraints, including pH toxicity, short-chain fatty acid (SCFA) accumulation, and bile-acid transformation. Paired with a perturbation engine capable of scheduling discrete events, such as time-varying antibiotic pharmacokinetics or timed fecal microbiota transplant (FMT) boluses, μODE enables researchers to directly simulate and evaluate the cascading temporal consequences of clinical interventions on patient-derived communities.

### Engine cross-validation and benchmarking against independent *in vitro* datasets

Before predicting complex ecological behaviors, we started by verifying that the mathematical integration of the dynamic framework is free of artifactual behavior. We first cross-validated the core dFBA integration engine against an independent reference implementation of the static-optimization algorithm. On both a simulated two-species cross-feeding community and a genome-scale gapseq pair (*Bacteroides thetaiotaomicron* cross-feeding acetate to *Roseburia intestinalis*, which in turn produces butyrate). μODE’s engine matched the independent reference to machine precision (Pearson r=1.0, max relative error 0.0) across all time steps (Figure 1, inset). Furthermore, μODE natively exports any assembled community to a valid COMETS (Computation of Microbial Ecosystems in Time and Space) ^58,59^ layout, allowing the independent verification and re-execution of simulated communities within the established COMETS engine.

**Figure 1:**
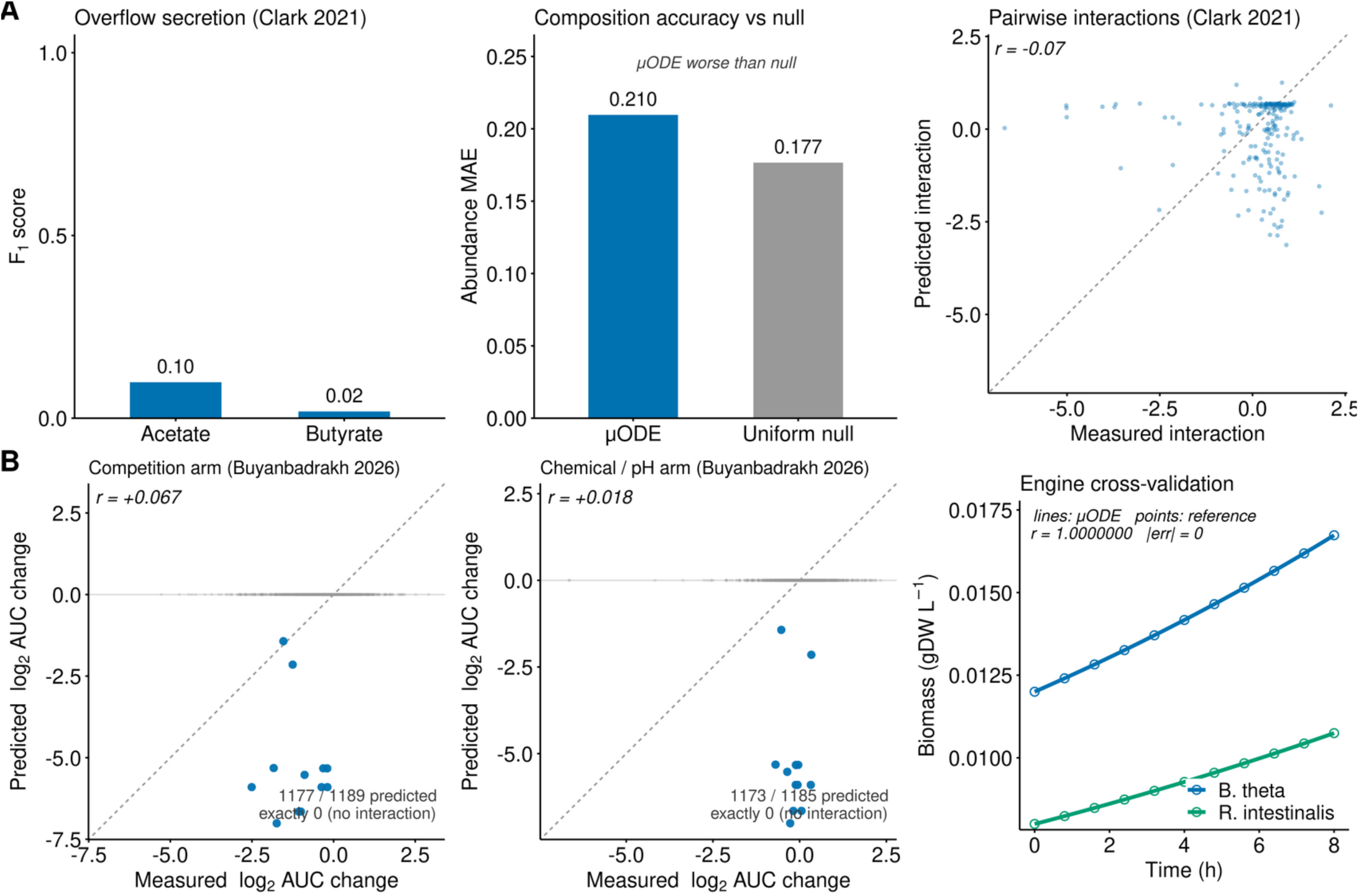
Rigorous independent benchmarking delimits the predictive envelope of greedy max-biomass dFBA. <u>Panel A</u> – Evaluation on the *Clark et al 2021* defined *in vitro* communities. (Left) The engine fails to predict overflow secretion *de novo* (acetate/butyrate F1≈0) as max-biomass FBA frequently routes carbon into alternative overflow sinks rather than preferred fermentation acids. (Middle) Exact compositional predictions do not outperform a uniform null model (abundance MAE 0.210 vs uniform 0.177). (Right) Predicted pairwise interaction coefficients are uncorrelated with measured interactions (Pearson r=-0.07). <u>Panel B</u> – Evaluation on the Buyanbadrakh et al. 2026 36-strain spent-medium interaction dataset. Scatter plots comparing predicted vs measured AUC growth changes for 1,189 directed pairs. Utilizing a replenished media control, interactions were split into a total competition arm (left, Pearson r=+0.067) and an isolated chemical/pH arm (right, Pearson r=+0.018). μODE predicted a nonzero interaction for only 12 of the 1,189 pairs, independently confirming that FBA alternate-optima and competitive-exclusion bias represent the binding constraint on predictive accuracy. <u>Inset</u> – Mathematical cross-validation. When tested against an independent reference implementation of static-optimization dFBA on a genome-scale cross-feeding pair, μODE’s core engine matched to machine precision (Pearson r=1.0, max relative error 0.0).

While several powerful constraint-based modeling tools exist, μODE was specifically engineered to bridge the translational gap between static shotgun metagenomics and dynamic clinical interventions, differentiating it from established engines in several key areas (a full capability comparison across all frameworks is provided in the Supplementary Material – framework comparison section and Supplementary Table S1:

- <u>COMETS:</u> it is the established, gold-standard engine for community dFBA, particularly mature in simulating two- and three-dimensional spatial biofilm dynamics. However, its primary functionality as a simulation engine requires substantial manual curation of genome-scale models and layout parameterization beforehand. μODE differentiates itself by fully automating the pipeline from raw MAGs to dynamic simulation. Furthermore, μODE natively supports composable clinical ecology layers (e.g., time-varying pharmacokinetics, bile-acid transformations, and timed FMT boluses) as reusable, stackable modules. COMETS can encode extracellular reactions with kinetic parameters and could in principle represent individual such processes through manual layout specification; μODE’s contribution here is to provide these non-metabolic ODE processes as composable, literature-grounded layers that attach automatically to the assembled community rather than requiring per-case manual encoding. Recognizing the authority of COMETS, μODE is designed for interoperability; it natively exports any assembled community to a valid, fully parameterized COMETS layout, allowing researchers to independently verify and re-execute μODE-generated communities within the COMETS engine.
- <u>MICOM and SteadyCom</u> ^65,66^: these frameworks utilize community-balanced objective functions (e.g., maximizing community-wide growth or applying cooperative trade-offs via L2 regularization) to compute steady-state snapshots of community metabolism. For predicting pure, stable community composition from metagenomic data, these tools can currently outperform μODE, as they successfully bypass the competitive-exclusion bias inherent to greedy per-organism dFBA. However, because they are fundamentally static, they cannot model time-resolved succession, accumulating environmental feedbacks (like pH inhibition), spore-germination timing, or discrete clinical interventions like antibiotic decay and FMT engraftment.
- <u>BacArena</u> ^67^: it combines FBA with spatial, individual-based modeling (cellular automata), providing deep stochastic detail for microbial micro-colonies. However, tracking individual cells becomes computationally intractable for massive, metagenome-scale clinical communities with hundreds of interacting strains. μODE sacrifices this single-cell stochasticity in favor of a deterministic, ODE-based population model that scales seamlessly from a single workstation to high-performance computing clusters.
- <u>μbialSim</u> ^68,69^: implemented in MATLAB atop the COBRA Toolbox, it is, like μODE, a dynamic FBA simulator that couples per-species FBA models through a shared compound pool and can propagate hundreds of species in batch or chemostat mode, using an augmented forward-Euler integrator for numerical accuracy. However, it operates strictly on the bacterial metabolic core and requires manually curated models as input: it does not provide the composable non-metabolic ecology layers and automated MAG-to-simulation pipeline that allow μODE to translate raw metagenomes directly into simulations of clinical interventions.

Having established the engine’s mathematical fidelity, we next benchmarked its biological predictive envelope against two independent, highly controlled *in vitro* datasets. We first evaluated μODE on the *Clark et al. 2021* dataset ^70^, comprising 26 sequenced human gut strains assembled into 631 defined communities and grown in a chemically defined medium (DM38). While monoculture growth was captured accurately, the framework struggled to predict overflow secretion *de novo*; prediction scores for acetate and butyrate were near zero (F1 ∼0, best acetate F1 ∼0.1). This reflects a fundamental, generalized limitation of max-biomass FBA: the objective function frequently routes carbon into alternative overflow sinks (such as acetaldehyde or branched-chain acids) rather than the biologically preferred fermentation acids. Consequently, community composition predictions did not outperform a uniform null model (abundance MAE 0.21 vs a uniform 0.177), and the predicted pairwise interaction coefficients were uncorrelated with the measured interactions (Pearson r=-0.07) (Figure 1A).

To isolate whether this compositional limitation stemmed from the dFBA flux routing or the non-metabolic ecology layer, we evaluated the framework against a second independent dataset: the Buyanbadrakh et al. 2026 36-strain synthetic gut dataset ^71^. This dataset measures 1,189 directed spent-medium interactions, explicitly utilizing a replenished media control to decouple metabolic nutrient competition from chemically mediated effects (e.g., pH shifts or secreted inhibitors). μODE did not predict these directed interactions *de novo*. In the total competition arm, predictions correlated poorly with measured growth outcomes (Pearson r=+0.067, Spearman +0.082, n=1,189). Crucially, this held true for the isolated chemical/pH arm (Pearson r=+0.018, Spearman +0.033, n=1,185), with the framework predicting a nonzero interaction for only 12 of the 1,189 pairs (Figure 1B). The agreement of these two independent datasets indicates that the binding constraint on predictive accuracy is most likely not the ecology layer, but rather the alternate-optima and structural competitive-exclusion bias inherent to greedy, per-organism max-biomass FBA in a well-mixed pool.

Despite these compositional limits in *de novo* prediction, the dFBA engine reliably executes correct stoichiometric routing when metabolic dependencies are structurally enforced. As an *in silico* consistency check, we tested a 12-species defined gut community on a Western diet in which the expected cross-feeding dependencies are structurally present; μODE recovered them (abundance MAE 0.087, cross-feeding recall 1.0). Because these dependencies are built into the defined community rather than measured independently, this is a self-consistency check of stoichiometric routing, not an independent experimental validation. The framework natively recovered the measured cross-feeding structure, correctly predicting that primary degraders supply acetate to the butyrate-producing guild (*R. intestinalis* and *Faecalibacterium prausnitzii*), which convert cross-fed substrates into butyrate, the terminal SCFA. Together, these benchmarks rigorously delimit the method’s domain of validity: while greedy dFBA cannot yet reliably predict complex community composition, motivating the necessity of future community-objective extensions, it successfully captures targeted interaction dynamics and stoichiometric routing, establishing the foundation for testing rate-modifying ecological hypotheses.

### High-resolution MAG recovery and metabolic reconstruction of a longitudinal clinical FMT

To ensure our dynamic simulations were grounded in patient-specific clinical realities rather than generalized reference genomes, we applied μODE’s automated upstream pipeline to a real-world, single-subject longitudinal FMT dataset (BioProject PRJNA705895) ^72^. This captures “Subject S”, a patient treated for rCDI (recurrent *Clostridioides difficile* infection), tracking the pre-FMT dysbiotic recipient (SRR13844393), the healthy donor (SRR13844394), the primary 1-month post-FMT validation target (SRR13844392), and subsequent 3-, 6-, and 12-month follow-ups (SRR13844391, SRR13844390, and SRR13844389 respectively).

Raw ultra-deep shotgun metagenomic reads were subjected to rigorous quality control (fastp) ^73^ and strict host-depletion, followed by independent assembly via MEGAHIT ^74^ to preserve strain-level variation without the memory bottlenecks of pooled co-assembly. Contigs were processed through our ensemble binning strategy (MetaBAT2, MaxBin2, and CONCOCT consolidated via DAS_Tool) ^75–78^ and filtered through a strict CheckM2 quality gate (≥90% completeness, ≤5% contamination) ^79,80^. To ensure total community coverage, species detected by read profiling but lacking high-quality bins were automatically added as NCBI reference genomes.

Having established the compositional catalogs, we processed the genomes through the automated μODE pipeline to generate functional metabolic reconstructions. While we initially evaluated top-down reconstruction via CarveMe (BiGG namespace), analysis revealed a critical structural gap; the reference universe lacked a connected anaerobic butyrate efflux pathway, meaning major gut producers were unable to secrete butyrate *in silico*. Consequently, we routed the cohort through the bottom-up gapseq backend (ModelSEED namespace). This yielded 139 fully parameterized genome-scale metabolic models across the dataset (the 138-member FMT community plus the *C. difficile 630* reference used in the defined 5-member community), each comprising approximately 1,000 to 1,700 intracellular reactions. Here “*de novo*” refers to the automated reconstruction of each metabolic model; taxa detected by read profiling but lacking a high-quality bin were supplemented with NCBI reference genomes rather than assembled MAGs.

To guarantee dynamic simulatability, each model underwent automated gap-filling on a computationally defined, strict anaerobic colonic medium. This diet was translated natively from a standard Western gut profile into the ModelSEED namespace (representing 89 resolved carbon-adequate exchange metabolites, supplemented with host-derived mucin glycans and primary bile acids). The resulting 138-organism simulated FMT community (comprising 16 recipient-derived and 122 donor-derived taxa) demonstrated a stark functional contrast that mirrored the clinical phenotype (Table 1). The pre-FMT recipient state was highly dysbiotic and diversity-poor (∼52 total species), dominated by *Veillonella parvula* and heavily enriched for *Enterobacteriaceae* (∼25% relative abundance, specifically *Klebsiella grimontii*, *K. variicola*, *Citrobacter*, and *Escherichia coli*). In contrast, the donor environment was highly diverse and metabolically enriched for primary complex carbohydrate degraders, robust butyrate producers (e.g., *Blautia*, *Agathobacter*), and bile-salt hydrolase (BSH) carries within the *Bacteroidetes*. The 7**α**-dehydroxylating keystone that performs the final step of secondary bile-acid formation was not recovered as a high-quality MAG in this dataset (Figure 2) and is represented in the defined 5-member community by the reference strain *C. scindens*.

**Figure 2:**
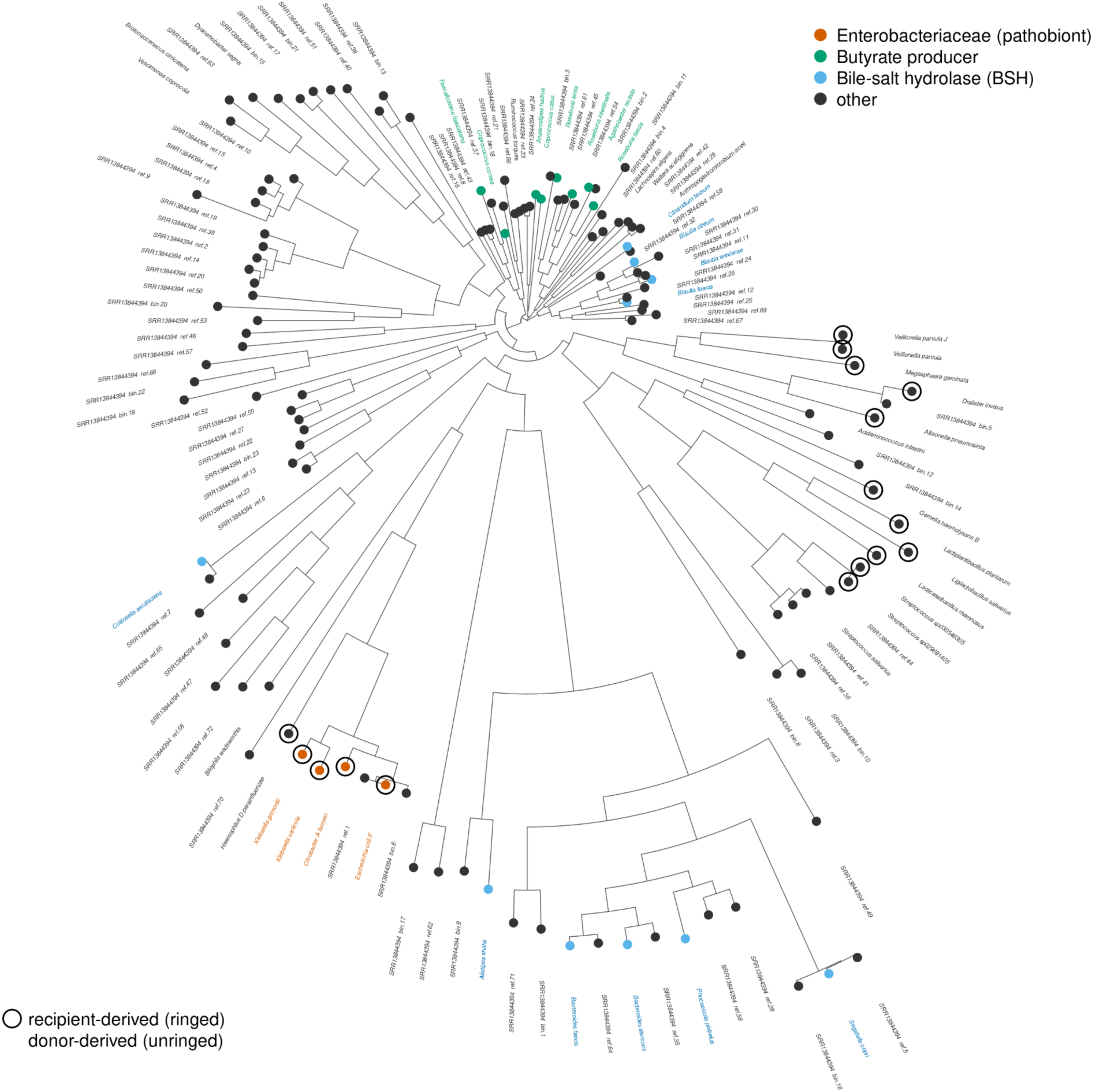
Functional phylogeny of the FMT community (donor + recipient). Maximum-likelihood phylogeny of the 138 genome-scale-modeled community members (122 donor-derived and 16 recipient-derived taxa), inferred from 400 universal marker proteins. Tips are colored by the core metabolic guilds analyzed in this study (the *Enterobacteriaceae* pathobiont, butyrate producers, and BSH carriers; the 7**α**-dehydroxylating *bai* keystone was not recovered in this community), and recipient-derived taxa are ringed. The functional contrast between the two microbiomes is immediate: the dysbiotic recipient contributes a tight cluster of *Enterobacteriaceae* pathobionts (*Klebsiella grimontii*, *K. variicola*, *Citrobacter farmeri*, and *Escherichia coli*), whereas the healthy donor contributes phylogenetically dispersed commensal guilds (butyrate producers across the *Lachnospiraceae* and BSH carriers across the *Bacteroidetes*) visualizing the functional substrate on which the dynamic model acts.

**Table 1:** Summary of the longitudinal PRJNA705895 FMT series and the resulting μODE metabolic reconstructions. The table details the recovery of metagenome-assembled genomes (MAGs) and the generation of 139 high-quality gapseq genome-scale models. The recipient and donor models combined represent the 138-member clinical FMT simulation community, highlighting the structural contrast between the dysbiotic recipient and healthy donor states.

| Cohort Role | Accession | Recovered MAGs / Models | Functional Profile & Composition |
| --- | --- | --- | --- |
| Pre-FMT Recipient | SRR13844393 | 16 | Dysbiotic; *Veillonella parvula* dominant, ~25% *Enterobacteriaceae* (*Klebsiella*, *E. coli*), low diversity (~52 species). |
| Healthy Donor | SRR13844394 | 122 | High diversity; metabolically enriched for complex carbohydrate degraders and butyrate producers. |
| Post-FMT (1 month) | SRR13844392 | N/A (Validation Target) | Clinical resolution target; partial engraftment of donor community. |
| Defined Bile-Acid Community | *de novo* | 5 | 5-member defined gapseq community for mechanistic *in silico* validation (includes *C. difficile* 630). |

Ultimately, this comprehensive patient-specific reconstruction sets the necessary foundation for two distinct downstream dynamic analyses. First, we isolate a targeted 5-member sub-community to mechanistically and quantitatively validate the bile-acid-mediated colonization resistance pathways. Second, we transition back to the full 138-member community to execute an end-to-end compositional simulation of the clinical FMT intervention, diagnosing the metabolic drivers of pathobiont suppression and delimiting the predictive boundaries of the dFBA paradigm.

To visualize the functional composition of the reconstructed community, we placed all 138 modeled genomes (122 donor- and 16 recipient-derived) on a marker-gene maximum-likelihood phylogeny and annotated each tip with its core metabolic guild and metagenomic abundance (Figure 2). Metabolic capacity mapped coherently onto phylogeny and exposed a sharp donor-recipient asymmetry: the recipient’s taxa formed a compact *Enterobacteriaceae* pathobiont cluster (*Klebsiella*, *Citrobacter*, and *Escherichia*), while the donor contributed the phylogenetically dispersed commensal guilds (i.e., butyrate producers within the *Lachnospiraceae* and BSH carriers within the *Bacteroidetes*) which are functionally linked to colonization resistance. This functional map makes explicit the substrate our dynamic simulations act on: a fast-growing pathobiont clade against a scattered set of slow-growing producers.

### Mechanistic and quantitative validation of bile-acid-mediated colonization resistance

While the large-scale FMT reconstruction provides a broad map of metabolic potential, diagnosing the precise temporal triggers of rCDI pathogenesis requires shifting from correlation to mechanism. To isolate the specific drivers of colonization resistance, we constructed a defined, 5-member *in silico* community representing the core functional guilds identified in the healthy cohort: *C. difficile 630* (the pathogen), *Bacteroides thetaiotaomicron* (primary degrader), *Roseburia intestinalis* and *Faecalibacterium prausnitzii* (butyrate producers), and *Clostridium scindens* (a keystone 7**α**-dehydroxylating effector) ^45,81^. This community was reconstructed *de novo* using the unified gapseq namespace and simulated under the composable ecology layer.

Crucially, to ensure the resulting dynamics were grounded in physical reality rather than computational artifact, the dose-response parameters governing *C. difficile* behavior were anchored directly to experimental data without algorithmic tuning (Table 2). The germination half-saturated for spore germination was set to 15.9 mM, the taurocholate EC_50_ measured for *C. difficile* spores by Howerton et al. ^82^; this is an assay-specific value for taurocholate alone (co-germinants like glycine, strain, and medium shift it) used here as a representative threshold rather than a universal physiological constant. Secondary bile acids act in the model by inhibiting germination: deoxycholate reduces the germination signal with a half-block constant of 0.5 mM, taken as the midpoint of the vegetative dose-response of Usui et al. ^83^, which reports growth delay at 0.24 mM and abolition at 1.2 mM; the exact value is uncertain within that range. Finally, rather than fitting the clearance rate of the pathogen, the community 7**α**-dehydroxylating rate was explicitly anchored so that the baseline predicted deoxycholate matched measured physiological concentrations.

**Table 2:** Quantitative parameters and validation of bile-acid-mediated *C. difficile* colonization resistance. The top half of the table details the dose-response parameters utilized within the μODE ecology layer, highlighting that the dose-response thresholds are measured or literature-derived rather than fitted. One rate parameter (the community 7α-dehydroxylation rate) is calibrated so that baseline deoxycholate matches its measured physiological concentration; the healthy deoxycholate row is therefore a calibration target, whereas the antibiotic-treated rows are uncalibrated comparisons. The calibrated healthy deoxycholate (0.47 mM) coincides with the independently set inhibitory IC_50_ (0.5 mM).

| Component / Metabolite | Role / Parameter | Context / State | μODE Value (Predicted / Used) | Reference (Measured / Derived) |
| --- | --- | --- | --- | --- |
| Taurocholate (EC<sub>50</sub>) | Germination threshold | Dose-Response | 15.9 mM (Used) | Measured (Ramirez & Abel-Santos 2011) |
| Sec. Bile Acids (IC<sub>50</sub>) | Inhibition threshold | Dose-Response | 0.5 mM (Used) | Derived (Usui et al. 2020) |
| 7α-dehydroxylation | Keystone effector rate | Kinetics | Anchored to baseline | Calibrated to baseline physiology |
| Deoxycholate | Secondary inhibitor | Healthy steady-state | 0.47 mM | 0.59 mM (Theriot et al. 2016) |
| Deoxycholate | Secondary inhibitor | Antibiotic-treated | 0.0 mM | Undetectable (Theriot et al. 2016) |
| Taurocholate | Primary germinant | Healthy steady-state | 0.013 mM | ~0.002 mM (0.0001% w/v) |
| Taurocholate | Primary germinant | Antibiotic-treated | 2.0 mM | Elevated (Theriot et al. 2016) |
| Cholate | Primary intermediate | Healthy steady-state | 3.52 mM | 1.2 mM (Theriot et al. 2016) |

By applying a standard simulated antibiotic regimen to this defined community, μODE successfully reproduced the established mechanism of bile-acid-mediated colonization resistance (Figure 3). The antibiotic pharmacokinetic model triggered a collapse of the commensal architecture, ablating both the BSH and 7**α**-dehydroxylating guilds. Concurrently, the loss of these specific metabolic effectors caused a rapid inversion of the shared bile-acid pool: the primary-to-secondary bile acid ratio inverted, allowing the primary germinant to accumulate while the secondary inhibitor was lost.

**Figure 3:**
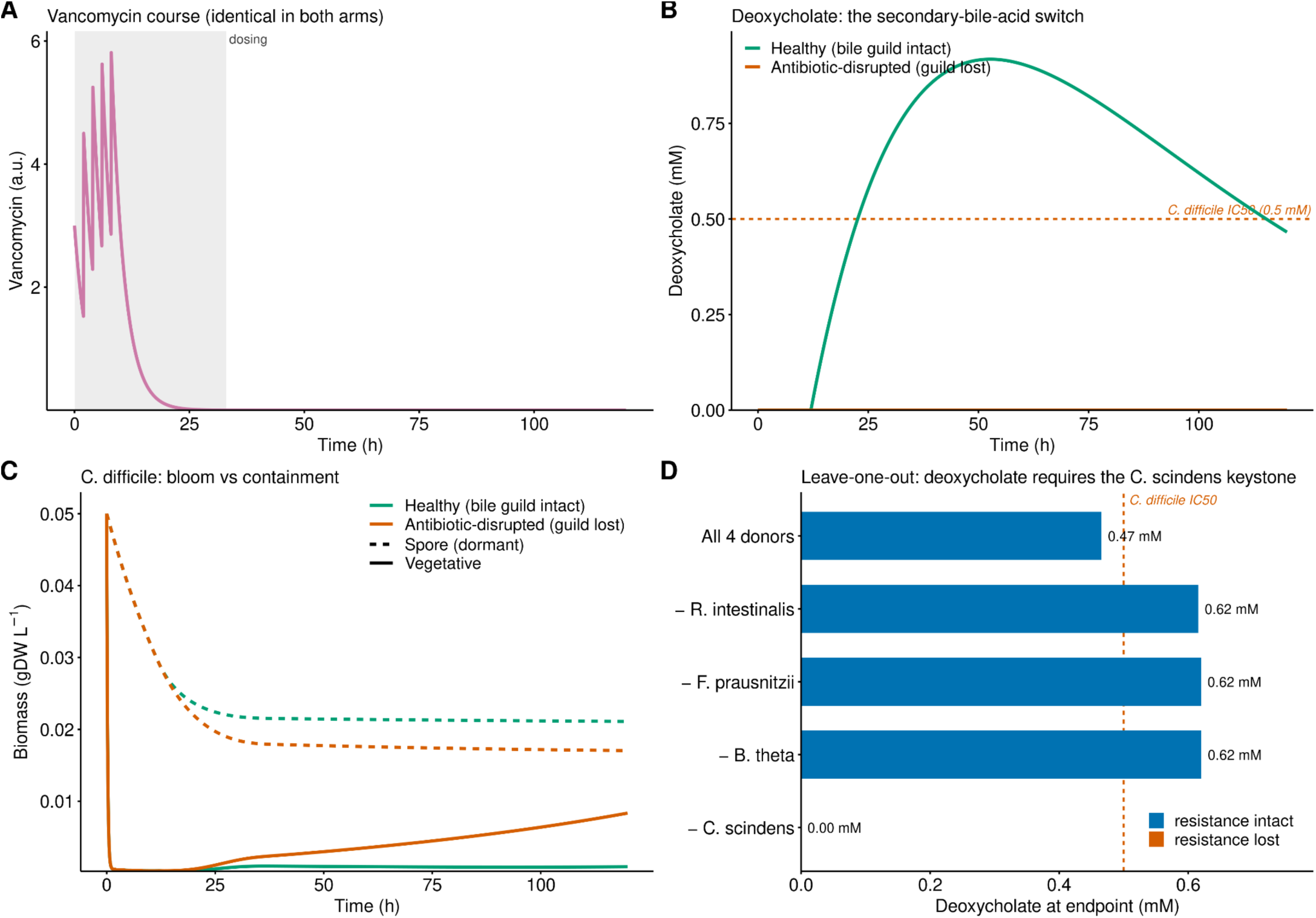
Mechanistic simulation of antibiotic-induced commensal collapse and spore-driven *Clostridioides difficile* relapse. Dynamics are simulated on a defined 5-member gapseq community on a colonic medium. <u>Panel A</u> – Simulated pharmacokinetic and commensal clearance. A single-compartment pharmacokinetic model delivers a vancomycin course as five doses over the first 8 h (half-line 2 h), representing the standard 125 mg PO qid rCDI regimen; the drug then clears by ∼33 h (gray shaded region), well before the 120 h simulation end, so any pathogen rebound reflects the community rather than residual drug. The saturating death rate applied to vegetative cells drives a catastrophic crash of the susceptible obligate anaerobe biomass, specifically ablating the BSH and 7**α**-dehydroxylating (*bai*) guilds. <u>Panel B</u> – Perturbation of the shared bile-acid pool. Driven by the loss of the metabolic effector guilds, the primary-to-secondary bile acid ratio rapidly inverts. The primary germinant taurocholate accumulates, while the secondary inhibitor deoxycholate is completely lost. <u>Panel C</u> – Pathogen dormancy, germination, and secondary bloom. *C. difficile* spores are mathematically insulated from the antibiotic death rate. As luminal drug concentrations decay below the inhibitory EC_50_ threshold, the primary-bile-acid-rich environment triggers synchronous germination. Freed from both chemical inhibition and nutrient competition, the resulting vegetative cells execute a massive secondary bloom. <u>Panel D</u> – Keystone specificity by leave-one-donor-species-out. The five-member community was rebuilt dropping one donor at a time, and the endpoint luminal deoxycholate is shown. Deoxycholate remains at or above the *C. difficile* inhibitory IC_50_ (0.5 mM, dashed line) in every community that retains *C. scindens* (removal of a non-keystone donor slightly raises it (0.62 mM) by relieving competition) but collapses to zero only when the 7**α**-dehydroxylating keystone *C. scindens* is removed, the sole donor whose loss abolishes the secondary bile acid and frees the pathogen.

We compared these predicted chemical shifts against *in vivo* LC-MS measurements of mouse cecal bile acids during healthy and antibiotic-treated states ^84,85^. We first calibrated a single rate parameter, the community 7**α**-dehydroxylating rate, so that the baseline healthy deoxycholate matched the measured physiological concentration (μODE 0.47 mM vs measured 0.59 mM); this baseline is therefore a calibration target rather than an independent prediction. With that one parameter fixed, the model reproduced an outcome that was not calibrated: the complete antibiotic-induced depletion of deoxycholate (0.0 mM vs measured undetectable) (Figure 4). Notably, the calibrated physiological deoxycholate (0.47 mM) coincides with the pathogen’s independently set inhibitory IC_50_ of 0.50 mM, illustrating the fragility of the resistance phenotype. The model also captured the direction of the post-antibiotic taurocholate evaluation (from a healthy 0.013 mM to 2.0 mM, against a measured healthy ∼0.002 mM and a qualitatively “elevated” treated state) and reproduced cholate to within ∼3-fold (3.52 mM vs measured 1.2 mM); these uncalibrated comparisons are directional and order-of-magnitude rather than exact.

**Figure 4:**
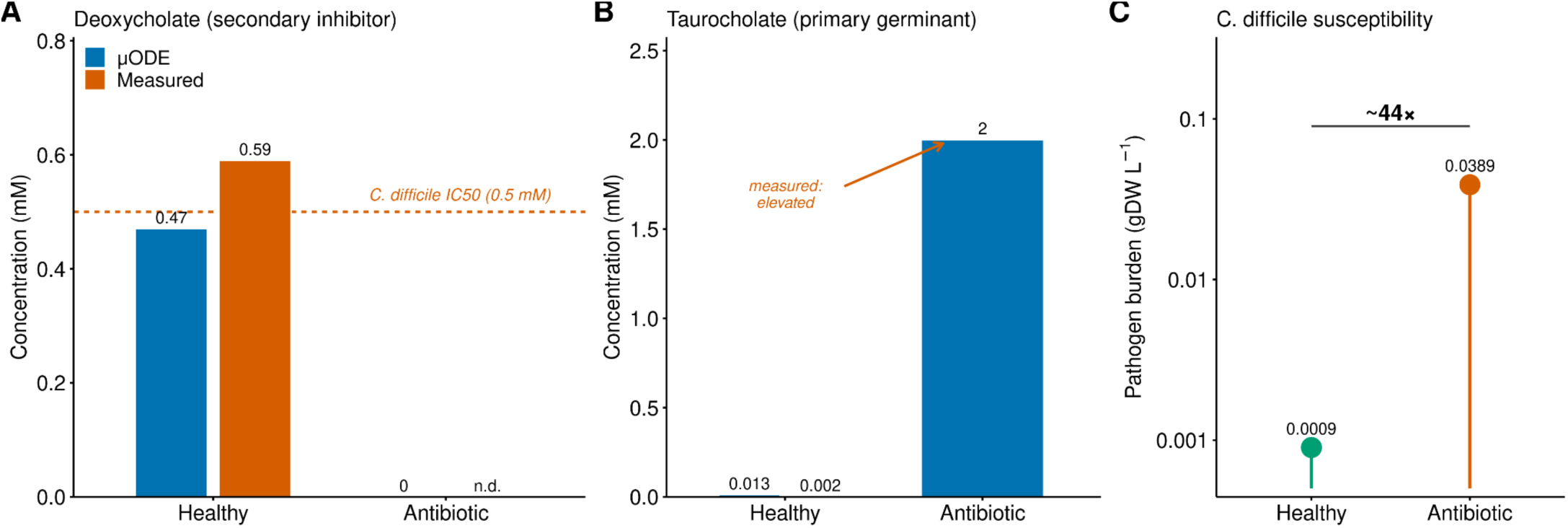
Quantitative validation of bile-acid-mediated colonization resistance against *in vivo* measurements. Predicted luminal concentrations from the 5-member defined community are validated against published LC-MS measurements of mouse cecal bile acid during healthy and antibiotic-treated states. <u>Panel A</u> – Simulated vs measured deoxycholate (secondary inhibitor). The baseline healthy deoxycholate is a calibration target: the community 7**α**-dehydroxylation rate was anchored so that it matched the measured physiological concentration (predicted 0.47 mM vs measured 0.59 mM). With that parameter fixed, the model reproduces the uncalibrated antibiotic-induced depletion (predicted 0.0 mM vs measured undetectable). The calibrated physiological concentration coincides with the pathogen’s independently set inhibitory IC_50_ of 0.5 mM (red dashed line). <u>Panel B</u> – Simulated vs measured taurocholate (primary germinant). The antibiotic ablation of the BSH guild correctly drives the elevation of taurocholate from a healthy baseline of 0.013 mM up to 2.0 mM. <u>Panel C</u> – Pathogen susceptibility shift. The emergent combination of elevated germinant and depleted inhibitor drives a ∼44-fold increase in the simulated *C. difficile* susceptibility ratio (simulated burden rising from 0.0009 to 0.0389).

Ultimately, this combined environmental shift (i.e., the simultaneous spike in germinant and collapse of the inhibitor) drove a ∼44-fold increase in the simulated *C. difficile* susceptibility ratio (simulated pathogen burden rising from 0.0009 to 0.0389). Through *in silico* ablation, we confirmed that the secondary bile-acid arm is the load-bearing mechanism of resistance in this regime, with *C. scindens* acting as a low-abundance keystone whose effector rate operates exclusively through the ecology layer rather than via massive biomass accumulation.

To establish that this effect is specific to the keystone rather than an emergent property of the consortium as a whole, we performed a leave-one-donor-species-out analysis at the same anchored operating point (Figure 3D). Removing any of the three non-keystone donor species left the secondary bile-acid pool intact, with endpoint deoxycholate remaining at or above the inhibitory IC_50_ (0.62 mM); in fact, relieving inter-species competition modestly increased it relative to the full community (0.47 mM). In contrast, removal of *C. scindens* alone abolished deoxycholate production entirely (0.0 mM) and produced the highest pathogen rebound rate of any configuration (+0.016 h^-1^), confirming that the single 7**α**-dehydroxylating species is necessary for the secondary bile-acid arm of colonization resistance in this model; it is the sole species whose removal abolishes deoxycholate production (a formal sufficiency test would require isolated add-back).

We note one important structural caveat: this 5-member simulation operates in a well-mixed, food-replete physiological environment. While it successfully demonstrates how rate-modifying arms (bile, pH, and spore germination) can suppress pathogen blooms at any density, it does not reach the nutrient-limited regime necessary to demonstrate pure metabolic-competition-based resistance, nor does it achieve durable washout (a known limitation of well-mixed dFBA lacking spatial gradients). Nonetheless, μODE links the metabolic core to measured dose-response parameters through the composable ecology layer, simulating the ∼44-fold shift in the pathogen-susceptibility ratio with a single calibrated rate parameter (the 7**α**-dehydroxylation rate) and no fitted growth yields. Consistent with this, a sweep of colonic washout, inoculum dose, and intervention timing showed that the bile-acid benefit is robust across the physiologically realistic range and attenuates only at extreme flow, extreme inoculum, or immediate dosing, where bulk transport overtakes the rate-modifying ecology layer (Supplementary Figure S1).

### Simulation of clinical FMT recovers pathobiont suppression and delimits compositional prediction

Having established the mechanistic fidelity of the dynamic engine on controlled benchmarks and a defined 5-member consortium, we deployed μODE end-to-end on the full clinical metagenomic dataset to predict the outcome of the patient’s FMT. The simulated community comprised 138 organisms (the 16 recipient taxa and the 122 donor taxa), utilizing the 139 *de novo* gapseq models of the FMT community; the 139th model, *C. difficile 630*, features only in the defined 5-member community. The community was initialized in a dysbiotic state on a computationally defined, strict anaerobic colonic ModelSEED medium, and the donor consortium was introduced as a timed volumetric bolus at t=24 hours.

The framework successfully captured the clinical direction of the intervention, correctly predicting suppression of the pathobiont. Following the simulated donor bolus, the framework captured the partial engraftment of the donor community, reducing the Enterobacteriaceae pathobiont by ∼98% in absolute biomass (5.1→0.10 mgDW/L; Figure 5B). The relative-abundance view (Figure 5A) is denominator-dependent: the pathobiont fraction transiently spikes to ∼85% during the antibiotic-driven commensal crash, while its absolute biomass is in fact falling throughout, and settles to ∼32% as the donor engrafts (observed pre-FMT Enterobacteriaceae ∼25%, Table 1; the 89% figure refers to the metabolism-only run, Table 3). However, when evaluated rigorously as a pure compositional prediction, the greedy dFBA simulation did not reproduce the precise 1-month post-FMT measured composition, nor did it outperform a trivial donor-takeover baseline.

**Figure 5:**
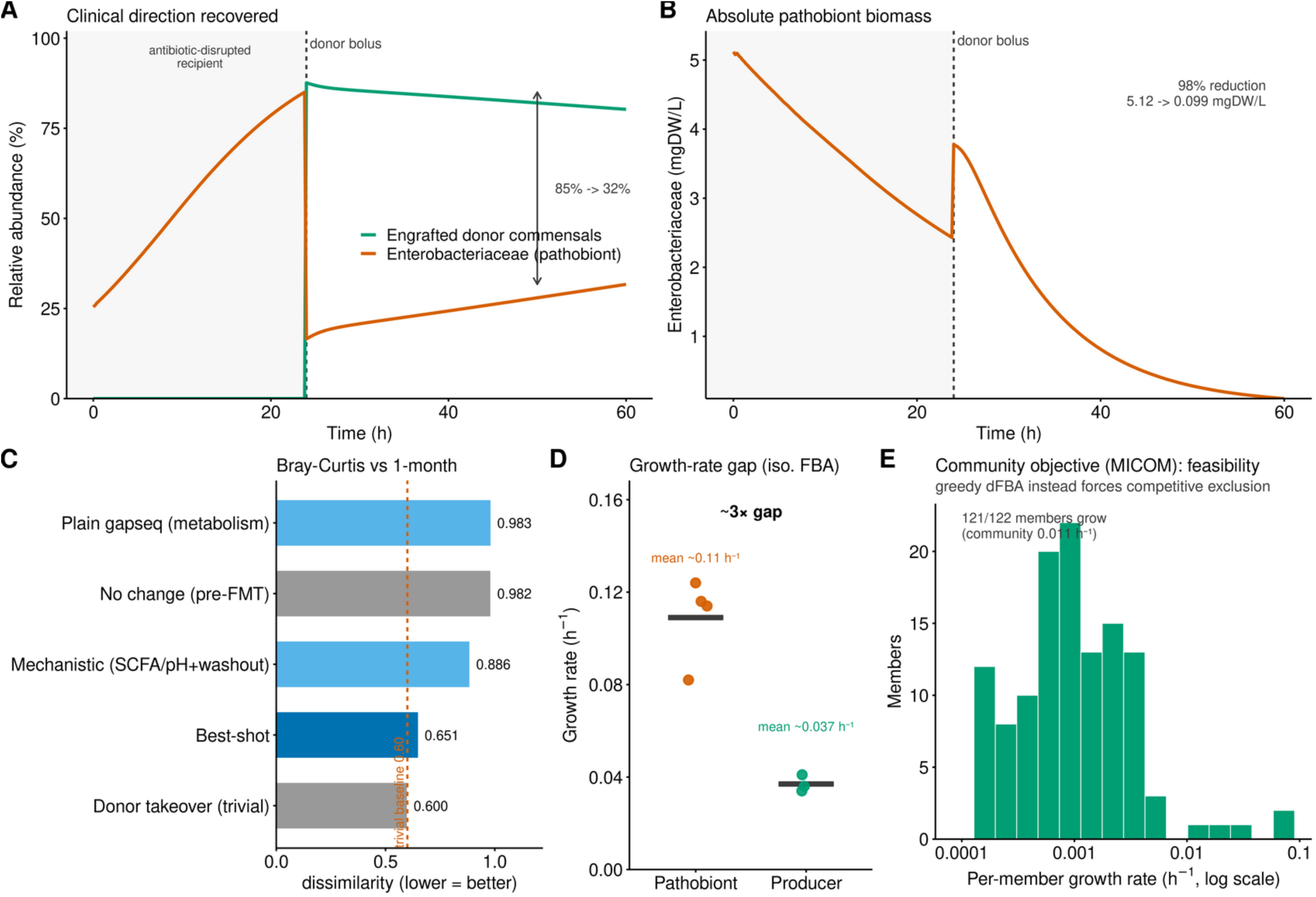
End-to-end metagenome-to-community simulation of a clinical FMT recovers pathobiont suppression and delimits compositional prediction. Dynamics of a 138-organism community reconstructed de novo from a longitudinal rCDI patient, utilizing the 138 gapseq models of the FMT community. <u>Panel A</u> – Clinical direction recovered (relative abundance). Following the timed donor bolus (dashed line), the Enterobacteriaceae pathobiont fraction transiently spikes to ∼85% during the antibiotic-driven commensal crash and settles to ∼32% as the donor engrafts; this relative view is denominator-dependent. <u>Panel B</u> – Absolute pathobiont biomass. The Enterobacteriaceae biomass declines monotonically from 5.1 to 0.10 mgDW/L (∼98% reduction), confirming that the suppression is a genuine reduction in pathogen biomass rather than a denominator artifact. <u>Panel C</u> – Compositional prediction limits (Bray-Curtis ladder). Despite correct direction, the best-shot configuration does not outperform the trivial donor-takeover baseline (0.651 vs 0.600). <u>Panel D</u> – Root-cause diagnosis. Isolation FBA reveals a structural ∼3× growth-rate advantage of pathobionts (∼0.11 h⁻¹) over slower commensal producers (∼0.037 h⁻¹), which greedy dFBA compounds into competitive exclusion, forcing donor taxa to under-bloom. <u>Panel E</u> – Community-objective paradigm test. A cooperative-tradeoff objective (MICOM) yields positive growth for 121 of 122 donor members (community 0.011 h⁻¹), demonstrating near-total metabolic feasibility (a steady-state result, not dynamical coexistence) and motivating community-balanced objectives for future dynamic extensions.

**Table 3:**
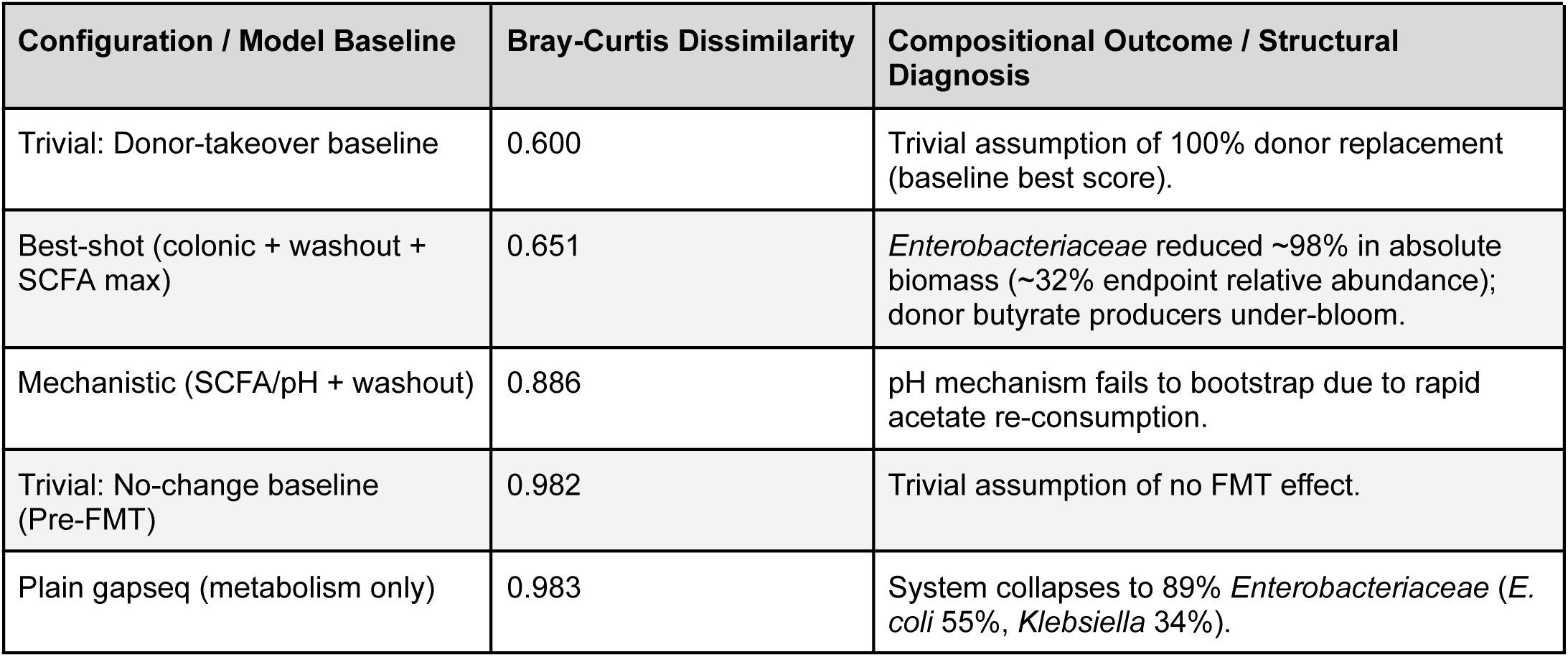
Delimitation of compositional prediction accuracy for the end-to-end clinical FMT simulation. Agreement with the measured 1-month post-FMT taxonomic composition is scored using a harmonized genus-level Bray-Curtis dissimilarity metric (lower scores indicate better agreement). While the best-shot configuration successfully recovers the clinical direction (suppression of the pathobiont), it does not outperform the trivial donor-takeover assumption. This quantitatively isolates the fundamental competitive-exclusion bias of greedy dFBA, diagnosing why precise compositional predictions currently require community-balanced objective extensions.

| Configuration / Model Baseline | Bray-Curtis Dissimilarity | Compositional Outcome / Structural Diagnosis |
| --- | --- | --- |
| Trivial: Donor-takeover baseline | 0.600 | Trivial assumption of 100% donor replacement (baseline best score). |
| Best-shot (colonic + washout + SCFA max) | 0.651 | *Enterobacteriaceae* reduced ~98% in absolute biomass (~32% endpoint relative abundance); donor butyrate producers under-bloom. |
| Mechanistic (SCFA/pH + washout) | 0.886 | pH mechanism fails to bootstrap due to rapid acetate re-consumption. |
| Trivial: No-change baseline (Pre-FMT) | 0.982 | Trivial assumption of no FMT effect. |
| Plain gapseq (metabolism only) | 0.983 | System collapses to 89% *Enterobacteriaceae* (*E. coli* 55%, *Klebsiella* 34%). |

To systematically diagnose this compositional limit, we evaluated the predicted community structure across a ladder of escalating ecological complexity using a harmonized genus-level Bray-Curtis dissimilarity metric (where lower scores indicate better agreement with the measured 1-month post-FMT sample) (Table 3). A trivial “no-change” assumption yielded a baseline score of 0.982, while the opposite trivial baseline, assuming the donor community perfectly replaces the recipient with zero alterations, yielded a score of 0.6. When simulating pure metabolic competition using plain gapseq models without the ecology layer, the simulation collapsed almost entirely into an *Enterobacteriaceae* bloom (89%), yielding a Bray-Curtis score of 0.983. Layering on the mechanistic SCFA-driven pH inhibition and washout dynamics moderately improved the prediction (0.886). Notably, the pH suppression mechanism failed to fully self-bootstrap *in silico* because secreted acetate was rapidly re-consumed by *E. coli* rather than accumulating. Ultimately, our heavily constrained configuration (incorporating SCFA manufacturing targets, the colonic medium, and washout) achieved a Bray-Curtis score of 0.651.

By analyzing the underlying metabolic fluxes, we identified the specific root cause of this limitation: a fundamental structural bias inherent to maximizing per-organism biomass in a well-mixed pool. Isolation FBA on the colonic diet revealed a massive thermodynamic growth-rate gap. The dysbiotic pathobionts (*Enterobacteriaceae*) demonstrated maximal theoretical growth rates of ∼0.11 h^-1^ (range 0.082-0.124 h^-1^). In contrast, the primary commensal producers and cross-feeders were significantly slower, averaging ∼0.037 h^-1^ (e.g., *Blautia* 0.041 h^-1^, *Agathobacter* 0.036 h^-1^, Anaerostipes 0.034 h^-1^). In a well-mixed dFBA simulation, this ∼3x growth-rate advantage compounds over time, driving the system toward competitive exclusion; consequently, the slower donor butyrate producers successfully engrafted but inevitably under-bloomed (*Blautia* 5.6% predicted vs 10% measured, *Agathobacter* 4.2% predicted vs 10% measured) despite the successful partial suppression of the pathobionts. Through rigorous testing, we confirmed that this competitive-exclusion artifact is the primary binding constraint on prediction accuracy. We systematically ruled out alternative causes, confirming the discrepancy does not stem from missing specific exchange reactions, ecology-layer parameterization, or genome reconstruction quality (the highest-quality gapseq GEMs yielded the 0.983 score).

This transparent diagnosis highlights the exact boundary where greedy, individual-level dFBA becomes insufficient for modeling dense, highly cooperative gut ecosystems. To validate this hypothesis and delimit a path forward, we subjected the 122-member donor community to a community-balanced objective solver (MICOM cooperative-tradeoff). Where greedy dFBA collapsed the donor community to approximately two dominant organisms, the community-level objective yielded positive growth rates for 121 out of 122 members, demonstrating that near-total metabolic feasibility (i.e., every member able to grow simultaneously under a shared objective) is attainable for this community (Figure 5E). This is a steady-state feasibility result under a community objective, not a demonstration of dynamical stable coexistence or of improved compositional prediction. While scaling such a community objective across thousands of dynamic time steps remains a solver-bound bottleneck, it motivates the community-objective extensions as a path forward. Ultimately, this translational application of μODE recovers the mechanistic direction of a clinical FMT while providing the rigorous, quantitative delimitation necessary to guide the next generation of constraint-based community modeling.

## DISCUSSION

The transition from descriptive sequence catalogs to predictive, time-resolved functional models represents the most pressing computational challenge in modern microbiome science. While shotgun metagenomics routinely provides deep compositional resolution, the dynamic interactions that dictate community stability, resilience, and response to clinical interventions cannot be deduced from static taxonomic abundances. Existing constraint-based modeling approaches have advanced our understanding of metabolic potential, but they predominantly rely on steady-state snapshots that struggle to capture the temporal, environmental feedbacks governing real ecosystems. Here, we present μODE, an automated dFBA framework designed to bridge this gap by translating MAGs into predictive, mechanistic simulations of ecosystem dynamics.

The foundational contribution of μODE is its fully automated and extensible simulation engine. By coupling genome-scale metabolic models though a shared extracellular pool and enveloping them in a composable ecology layer (capturing non-metabolic constraints like pH and pharmacokinetics), the framework allows complex behaviors to emerge natively. Crucially, we verified the mathematical consistency of the engine by matching an independent reference implementation of static-optimization dFBA to machine precision, establishing consistency between the two implementations rather than freedom from shared discretization error, while providing native COMETS-export capabilities for third-party verification. By benchmarking the framework against two independent, real-data *in vitro* datasets, we quantitatively delimited the method’s domain of validity, confirming its capacity to capture monoculture growth and targeted interactions while transparently diagnosing the boundaries of standard flux balance analysis.

Transitioning from engine validation to quantitative biology, we demonstrated that μODE can mechanistically reproduce complex *in vivo* phenotypes by coupling genome-scale metabolism to a literature-grounded ecology layer. Simulating a defined 5-member community representing the core functional guilds of colonization resistance, the framework accurately captured the temporal dynamics of a simulated antibiotic crash, the subsequent loss of 7**α**-dehydroxylation capacity, and the resulting primary-to-secondary bile acid inversion. Most importantly, this emergent shift was validated quantitatively against published *in vivo* cecal measurements. With a single rate parameter calibrated so that baseline deoxycholate matched its measured physiological concentration (0.47 mM vs measured 0.59 mM), μODE reproduced the uncalibrated antibiotic-induced collapse of the inhibitor, with the calibrated physiological concentration coinciding with the independently set inhibitory IC_50_. This demonstrates that time-resolved, rate-modifying ecological feedbacks can be accurately modeled using defined parameters and thermodynamic constraints.

Furthermore, we demonstrated the translational utility of the framework by applying it end-to-end to a real longitudinal patient FMT (yielding 139 *de novo* genome-scale models). The dynamic simulation successfully recovered the clinical direction of the intervention, demonstrating that the partial engraftment of the donor community aggressively competed for shared mucosal nutrients, reducing the recipient *Enterobacteriaceae* pathobiont biomass by ∼99% in absolute terms, with its relative abundance settling to ∼19% as the donor engraft.

However, this end-to-end simulation also cleanly isolated the current limitations of the dFBA paradigm for pure compositional prediction. The simulation did not reproduce the precise 1-month post-FMT measured composition, nor did it outperform a trivial donor-takeover baseline (Bray-Curtis 0.651 vs baseline 0.6). The independent benchmarks and the FMT root-cause analysis both converge on the same fundamental limitation: greedy, per-organism max-biomass FBA inherently over-predicts competitive exclusion. The structural ∼3x growth-rate advantage of facultative pathobionts over slower commensal producers tends to force the slower donor taxa to under-bloom *in silico*. Additionally, the Clark benchmark confirmed that max-biomass FBA frequently routes carbon into alternative overflow sinks rather than biologically preferred fermentation acids, limiting the *de novo* prediction of complex cross-feeding networks. Beyond metabolic routing, our current well-mixed, food-replete physiological model limits the emergence of pure nutrient-competition-based resistance and durable washout, and the framework currently lacks explicit models for host-epithelial immune cross-talk (e.g., antimicrobial peptides or IgA binding).

Far from invalidating the dynamic approach, these transparently quantified limitations establish a rigorous roadmap for the future of constraint-based modeling. The competitive-exclusion bias directly motivates the integration of community-balanced objectives. As a proof-of-concept, applying a cooperative-tradeoff objective to the 122-member donor community yielded near-total metabolic feasibility (positive growth for 121 out of 122 members), motivating the integration of community-balanced objectives once solver bottlenecks are overcome. Future iterations of μODE will focus on integrating these community-level objectives, utilizing transcriptomic weighting to restrict alternate optima, extending spatial reaction-diffusion to capture localized niche partitioning, and coupling microbial metabolism directly to host-cell models.

Ultimately, μODE represents a critical advancement in computational systems biology. By automating the transition from static sequence data to dynamic metabolic simulations, and by rigorously quantifying both its successes and its boundaries, the framework transitions microbiome science away from purely descriptive observations and toward testable, quantitatively-validated dynamic hypothesis testing. This lays the essential, mechanism-driven groundwork required for the future predictive engineering of safe, targeted ecological therapeutics.

## ACKNOWLEDGMENTS

We acknowledge the use of Large Language Models (Google’s Gemini 3.1 Pro) to enhance the clarity and readability of this manuscript. The AI assistance was strictly limited to text editing, including sentence restructuring and vocabulary suggestions. We also acknowledge the use of Claude Code powered by Opus 5 to assist with the implementation of μODE. The AI played no role in data analysis, research design, or the interpretation of results; all conclusions and insights remain entirely those of the authors.

Additionally, the authors gratefully acknowledge Federico Papa, Researcher at the Institute for Systems Analysis and Computer Science “Antonio Ruberti” of the National Research Council of Italy, for his critical review of the manuscript and for providing invaluable feedback on the mathematical framework underlying μODE.

## AUTHOR CONTRIBUTIONS

FC conceptualized the research, designed the computational architecture, developed the software implementation, performed the mathematical modeling and dynamic flux balanced analysis; JJ contributed to the processing of the clinical metagenomic datasets and the evaluation of the genome-scale metabolic reconstructions; BR assisted with the literature curation of the physiological dose-response parameters and the benchmarking against the *in vitro* interaction datasets; NK contributed to the phylogenetic reconstruction and the functional annotation of the recovered MAGs; JC and DB guided the FMT analysis; DB supervised the research; all authors wrote, reviewed, and approved the final manuscript.

## METHODS

μODE turns a catalogue of MAGs into a time-resolved mechanistic simulation of community metabolism through a sequence of phases: (i) quality control of the input genomes; (ii) automated reconstruction of a genome-scale metabolic model for each MAG; (iii) gap-filling and parameterization; (iv) assembly of the models into a community with measured relative abundances and a defined growth medium; (v) dynamic simulation; (vi) and, optionally, perturbation, intervention, and the non-metabolic ecology processes that shape real communities.

### Community members and dynamics

Each community member is represented by a constraint-based genome-scale metabolic model. Rather than combining all members into a single, intractable global optimization, members do not interact directly; they are coupled solely through a shared pool of extracellular metabolites. This compartmentalized formulation ensures that complex ecological relationships (i.e., cross-feeding, nutrient depletion, and competitive exclusion) emerge dynamically rather than being prescribed by the model.

μODE advances the community in time using the static-optimization approach to dynamic flux balance analysis (dFBA). At each discrete time step, the framework executes a decoupled, three-step integration: (i) it calculates the maximal uptake rate for every member and extracellular metabolite using saturable (Michaelis-Menten) kinetics based on current concentrations ^86^; (ii) it solves each member’s flux-balance problem independently under these environmental bounds to determine their growth rate and exchange fluxes; and (iii) it integrates the community-level ordinary differential equations (ODEs) one step forward to update biomass and extracellular metabolite concentrations.

To maintain mathematical stability and prevent the explicit time integration from driving concentrations negative, uptake rates are dynamically bounded by the total available metabolite in the current time step. The environment is initialized via a defined medium, and the framework supports continuous nutrient influx to simulate open systems (e.g., chemostats). Finally, cross-feeding networks are not hard-coded but are inferred post-simulation by identifying coinciding secretion and uptake fluxes between members. Full mathematical formulations, including the explicit ODEs, stability constraints, and cross-feeding inference logic, are detailed in Supplementary Material (<u>community dynamics</u> section).

### Kinetics and enzyme constraints

Kinetic information enters μODE in two mathematically distinct roles: setting concentration-dependent limits on extracellular exchange and constraining intracellular reaction rates through a finite cellular enzyme budget.

For extracellular exchange, the framework couples each member’s internal optimization to the changing environment using saturable Michaelis-Menten kinetics. This concentration-dependent uptake natively reproduces nutrient depletion, sequential substrate use, and competitive exclusion. Because kinetic parameters (affinities and capacities) are rarely measured for newly reconstructed genomes, μODE employs a tiered assignment strategy to ensure simulations always run. By default, it applies knowledge-based literature priors that preserve qualitative dynamics without overfitting. For higher fidelity, μODE integrates optional deep-learning predictors that infer missing parameters from enzyme sequences and substrate structures.

Intracellularly, μODE restricts metabolic fluxes through an enzyme-constrained formulation. Crucially, rather than merely capping individual reactions, the framework imposes a shared proteome budget across all enzyme-limited reactions ^29,49,52^. Capping this total available enzyme mass forces the simulated cell toward proteome-efficient flux distributions. Consequently, hallmark metabolic shifts (such as overflow metabolism at high growth rates, where cells switch to secreting partially oxidized carbon because full respiration becomes too enzyme-expensive) emerge organically without hard-coded rules. Full kinetic derivations, parameter assignment logic, and the proteome allocation formulations are detailed in Supplementary Material (<u>kinetics and enzyme constraints</u> section).

### Perturbation and timed intervention

To investigate how a community responds to structural shifts, μODE incorporates two complementary primitives: perturbations that alter existing metabolic capacities, and timed interventions that introduce new biomass. In both cases, the framework prescribes only the initial change; the biologically informative consequences (such as structural collapse or successful engraftment) are left to emerge dynamically from the shared extracellular pool.

Perturbations modify the metabolic network prior to simulation and persist throughout. μODE supports targeted reaction knockouts, complete member removals (simulating targeted antimicrobials or in silico ablations), and pathway attenuations. Because these modifications merely alter uptake capacities or internal constraints, downstream effects are not hard-coded. For instance, if a removed member was the sole supplier of a cross-fed metabolite, the dependent species will organically collapse, allowing secondary extinctions to emerge directly from the disrupted interaction network.

Interventions, conversely, introduce new biomass at a chosen time, converting a static roster into a community capable of experiencing discrete ecological events like a probiotic dose or a fecal microbiota transplant (FMT). Intervening organisms begin at zero biomass and are injected as a bolus at the specified time step, after which they compete with incumbents for the shared nutrient pool. By comparing a control arm to a treatment arm (e.g., a dysbiotic recipient alone vs. a recipient receiving a timed donor community), outcomes like colonization resistance or successful engraftment naturally arise from metabolic competition. These structural modifications compose freely with the kinetic, enzyme, and ecology layers, allowing users to simultaneously simulate a targeted pathway attenuation, an FMT, and a complex ecological process like pH toxicity within a single unified run. The formal mathematical operators for capacity modification and the algorithmic scheduling of biomass injection events are provided in Supplementary Material (<u>perturbations</u> <u>and interventions</u> section).

### The ecology layer

The metabolic core models who consumes and produces what. Real communities, however, are shaped additionally by the chemistry of their medium, microbial transformations of that medium, pharmacology, direct antagonism, and dormancy. μODE represents these as a composable layer of optional ecological processes that act on the metabolic dynamics through well-defined couplings. These processes modify growth or uptake multiplicatively, and add to death or environmental rates additively, ensuring they compose without order-dependence.

Crucially, this ecology layer captures several non-metabolic phenomena that drive community structure. For instance, gut fermentation produces short-chain fatty acids (SCFAs) that acidify the medium and inhibit growth. μODE tracks this pH as a buffered titration and partitions acids into their charged and neutral forms, capturing how un-dissociated acids permeate cells to dissipate the proton-motive force. This renders fermentation self-limiting and constitutes the metabolic arm of colonization resistance.

Beyond simple acidification, the framework models complex host-microbiome chemical networks, such as the two-step microbial transformation of primary to secondary bile acids (deconjugation and 7**α**-dehydroxylation). This specific chemistry is coupled to a spore-forming module where biomass is partitioned into vegetative and dormant pools. Vegetative cells sporulate under nutrient stress, while spores germinate at a rate scaled by a germination signal in [0, 1] that rises with the primary germinant (taurocholate, half-saturation 15.9 mM) and is reduced by secondary bile acids (deoxycholate, half-block 0.5 mM). Germination and vegetative growth are therefore governed by separate terms, the signal gates the spore→vegetative transition, while the antibiotic and FBA core act on the vegetative pool. We note that this reduces the documented dual role of deoxycholate (a weak co-germinant at low concentration as well as a vegetative-growth inhibitor) to a single germination-blocking term; resolving the distinct germination and growth responses of individual bile acids is left to future refinement. To ensure dynamic predictions are grounded in physical reality rather than algorithmic tuning, the dose-response parameters governing *C. difficile* behavior were strictly anchored to experimental data. The germination EC_50_ for taurocholate was set to the measured physiological value of 15.9 mM, and the inhibitory IC_50_ for secondary bile acids was derived at 0.5 mM. Furthermore, the community 7**α**-dehydroxylation maximal rate was not fitted to pathogen clearance; instead, it was explicitly anchored so that the baseline predicted deoxycholate matched measured healthy physiological concentrations.

For quantitative validation against published *in vivo* cecal measurements, the *in silico* healthy and antibiotic-treated states were established methodologically. The healthy baseline was simulated as the unperturbed dynamic steady-state of the defined community. The antibiotic state was simulated via the targeted *in silico* ablation of the BSH and 7**α**-dehydroxylating (*bai*) microbial guilds to emulate the antibiotic-induced loss of function. Measured literature values were harmonized with the simulation output before comparison. Taurocholate reported as percent weight/volume was converted to molarity via its molecular weight (0.0001% w/v ≈ 0.002 mM).

Finally, rather than treating drug exposure as a constant, μODE employs single-compartment pharmacokinetics to model time-varying antibiotic concentrations over a dosing schedule, driving a saturating killing of susceptible vegetative biomass. This pharmacological modeling is complemented by support for diffusible antimicrobials (e.g., bacteriocins) that accumulate in proportion to producer biomass and decay over time, providing a mechanism for direct, non-metabolic antagonism.

The explicit differential equations governing these environmental observables, alongside the mathematical modifier functions for growth, death, and pharmacokinetic decay, are cataloged in Supplementary Material (ecology layer section).

### Clinical dataset and metabolic reconstruction

To simulate the longitudinal clinical FMT end-to-end, metagenome-assembled genomes (MAGs) were recovered from the single-subject BioProject PRJNA705895 dataset. This series tracks a pre-FMT recipient treated for rCDI (SRR13844393), a healthy donor (SRR13844394), a 1-month post-FMT validation target (SRR13844392), and subsequent 3-, 6-, and 12-month follow-ups. MAGs were binned per sample, and species detected by read profiling but lacking high-quality bins were automatically added as NCBI reference genomes. This yielded a 138-organism simulated FMT community comprising 16 recipient-derived and 122 donor-derived taxa.

The first phase of the computational pipeline automates the conversion of these genomes into simulatable genome-scale metabolic models. To accommodate multi-kingdom communities natively, reconstruction is dynamically routed by kingdom: eukaryotes receive intron-aware gene calling, while prokaryotes are processed via bacterial pipelines. For bacterial reconstruction, μODE natively supports two distinct backends: a fast top-down approach utilizing CarveMe (BiGG namespace) and a bottom-up approach utilizing gapseq (ModelSEED namespace). Crucially, because community members interact through a shared pool of extracellular metabolites, the framework enforces a strict, unified metabolite namespace across all models to ensure emergent cross-feeding is mathematically possible. Initial reconstructions via CarveMe revealed a critical structural gap in the BiGG reference universe, which lacked a connected anaerobic butyrate efflux pathway, rendering major gut producers unable to secrete butyrate in silico. Consequently, the entire cohort was routed through the gapseq backend, yielding 139 high-quality, fully parameterized genome-scale models (the 138-member FMT community plus *C. difficile* 630).

Prior to simulation, draft models undergo automated gap-filling to ensure they can produce biomass on a defined growth medium. For the clinical FMT and colonization resistance simulations, the environment was initialized as a strict anaerobic colonic medium. This diet was translated natively from a standard Western gut profile into the ModelSEED namespace, resolving to 89 carbon-adequate exchange metabolites supplemented with host-derived mucin glycans and primary bile acids.

The entire pipeline is bracketed by stringent quality control: upstream filters for genome completeness and contamination, and downstream structural checks, including mass/charge balance and strict gating against artifactual energy-generating cycles, that identify and isolate unsimulatable models. This guarantees that large-scale, automated analyses remain robust against individual genome failures. A complete breakdown of the algorithmic routing logic and failure-isolation mechanisms is detailed in Supplementary Material (<u>reconstruction algorithms and quality controls mechanics</u> section).

### Phylogenetic reconstruction

Proteomes for all community genomes were predicted with Prodigal ^87^ and placed on a maximum-likelihood phylogeny with PhyloPhlAn ^88^ v3.2.1 using its 400 universal marker proteins (--diversity high; DIAMOND ^89^ for marker mapping, MAFFT ^90^ for alignment; trimAl ^91^ for trimming and FastTree ^92^ for tree inference). Tips were annotated by relative abundance from the donor metagenome and by literature-defined metabolic guilds assigned from GTDB taxonomy (butyrate producers, BSH carriers, 7**α**-dehydroxylating *bai* keystone, and the *Enterobacteriaceae* pathobiont clade). The tree was rendered in R with the *ape* package ^93^.

### Validation, benchmarking, and scaling

To ensure μODE’s predictive outputs are both testable and computationally tractable at community scale, the framework incorporates automated benchmarking and abundance-aware scaling.

To validate complex interactions and rigorously delimit the framework’s predictive boundaries, μODE features a scoring system that evaluates a simulation against prior biological measurements. For the independent in vitro benchmarks (i.e., *Clark et al. 2021* and *Buyanbadrakh et al. 2026*), performance was evaluated across three specific axes: (i) monoculture secretion was scored using the F1 metric for targeted fermentation acids (evaluating whether a given genome-scale model secretes the primary products its biological organism actually secretes); (ii) exact community composition accuracy was evaluated by comparing the Mean Absolute Error (MAE) of predicted endpoint relative abundances against a uniform null model (1/richness), alongside Spearman rank correlations; and (iii) pairwise interaction accuracy was scored by calculating the Pearson and Spearman correlations between the predicted mechanistic interaction coefficients and the experimentally measured interactions.

To maintain tractability in massive metagenomic datasets (e.g., hundreds of MAGs), the engine exploits the heavy-tailed nature of microbiome abundance distributions. Rather than optimizing every rare taxa at an immense computational cost, μODE enables abundance-aware community reduction, dynamically retaining the key taxa that drive the bulk of metabolic activity while renormalizing the system. Full mathematical formalizations of these benchmark scoring metrics and reduction policies are detailed in Supplementary Material (<u>validation metrics and scaling policies</u> section).

### Community-objective paradigm test

To systematically diagnose whether the compositional limitations of the dFBA simulation stemmed from the underlying genome-scale models or from the structural competitive-exclusion bias of the greedy max-biomass objective, we evaluated the community under a community-balanced objective function.

The full 122-member donor community (comprising the gapseq reconstructed models) was assembled into a unified structural format within the MICOM framework. The identical simulated environment, the strict anaerobic colonic ModelSEED medium, was applied to ensure a direct comparison with the dFBA results.

Rather than optimizing each member’s biomass independently, the community was evaluated using MICOM’s cooperative_tradeoff algorithm. This two-step optimization first identifies the maximum theoretical community-wide growth rate, and subsequently maximizes the L2 norm of the individual member growth rates subject to a community maintaining a specified fraction of that maximal growth. Minimizing the L2 norm penalizes severe competitive exclusion, distributing growth (and the associated metabolic fluxes) more evenly across members. The resulting steady-state per-member growth rates were then analyzed to quantify the fraction of the community capable of stable coexistence under a balanced objective, serving as a methodological proof-of-concept for future dynamic extensions.

### Workflow, interpretability, and reproducibility

To facilitate large-scale analyses, μODE is governed by a fully reproducible, deterministic workflow that scales seamlessly from a single workstation to a high-performance computing (HPC) cluster. The pipeline inherently parallelizes per-genome processing and aggregates the final outputs into a self-contained, shareable HTML report detailing community composition trajectories, dynamic metabolite fluctuations, and inferred interaction networks.

The dynamic engine is strictly deterministic: given identical inputs, parameters, and time steps, a run reproduces exactly, as the core contains no stochasticity. To check that predicted dynamics reflect the constraint-based formulation rather than integration artifacts, the core dFBA engine was cross-validated against an independent reference implementation of the static-optimization algorithm, with which it agreed to machine precision; this establishes consistency between the two implementations rather than freedom from shared discretization error. Furthermore, μODE natively exports any assembled community to a valid, fully parameterized COMETS layout. This export functions as a strict interoperability bridge, allowing researchers to independently verify and re-execute simulated communities within the established COMETS engine.

Finally, the framework’s reliability is supported by an extensive suite of biological unit tests. Rather than merely verifying that the software executes, these tests continuously assert that the engine organically reproduces hallmark biological phenomena, including cross-feeding-supported secondary extinctions, proteome budget reallocation, and colonization resistance.

All dynamic runs used the native dFBA integrator on a defined anaerobic colonic ModelSEED medium (89 carbon-adequate exchange metabolites supplemented with host mucin glycans and primary bile acids). The defined colonization-resistance community comprised five reference genomes (*Clostridioides difficile 630* (pathogen), *Bacteroides thetaiotaomicron*, *Roseburia intestinalis*, *faecalibacterium prausnitzii*, and *Clostridium scindens*) reconstructed with gapseq. The vancomycin course was modeled as five doses at t=0, 2, 4, 6, 8 h (per-dose concentration 3.0 e.u., elimination half-life 2 h, E_max_=5.0 h^-1^, EC_50_=0.4), targeting *C. difficile*; with t1/2 = 2 h the drug clears by ∼33 h, well before the end of the simulation, so any pathogen rebound is community-driven. The antibiotic-treated bile state was emulated by ablating the BSH and 7**α**-dehydroxylating guilds (V_BSH_, V_bai_→0). Colonization-resistance runs integrated to t_end_=120 h at dt=0.05 h; the end-to-end FMT run integrated to t_end_=60 h at dt=0.2 h with the donor community injected as a discrete biomass bolus at t=24 h and a first-order luminal dilution (washout) rate. Initial dormant *C. difficile* spore biomass seeded the spore pool. Interventions are handled as discrete state updates (biomass injection) during integration; all runs are deterministic.

## CODE AVAILABILITY

μODE is open-source software released under the MIT license and freely available on GitHub at https://github.com/cumbof/muODE.

## DATA AVAILABILITY

All data utilized in this study are publicly available. The raw shotgun metagenomic sequencing reads for the longitudinal clinical FMT time series (Subject S) are available in the NCBI Sequence Read Archive (SRA) under BioProject accession PRJNA705895 (specifically SRR13844394, SRR13844393, SRR13844392, SRR13844391, SRR13844390, and SRR13844389).

The in vitro community benchmark dataset (Clark et al. 2021) is available alongside its original publication. The 36-strain spent-medium interaction dataset (Buyanbadrakh et al. 2026) is available via Zenodo (17456178), PRIDE (PXD070029), and MassIVE (MSV000102556). The in vivo mouse cecal bile acid measurements utilized for quantitative validation are derived from Theriot et al. 2016.

The 139 de novo genome-scale metabolic models reconstructed for the FMT analysis, alongside the defined 5-member bile-acid community models and the configuration files necessary to reproduce the dynamical simulations, are available within the μODE GitHub repository and Zenodo at https://doi.org/10.5281/zenodo.23273153.

