## Supplementary Material for "Mechanistic prediction of fecal microbiota transplant engraftment and *Clostridioides difficile* colonization resistance with μODE"

**FRAMEWORK COMPARISON**

Established constrain-based community tools occupy distinct corners of the design space (Supplementary Table S1): COMETS is the mature dynamic, spatially explicit dFBA engine; MICOM and SteadyCom compute steady-state community optima under a community-level objective; BacArena resolves stochastic single-cell agents; and µbialSim provides an accurate well-mixed dFBA integrator. On the metabolic core these engines share, µODE is numerically faithful: its static-optimization integrator reproduces an independent reference dFBA implementation to machine precision (Pearson $r$ = 1.0) and exports any assembled community to a runnable COMETS layout for third-party verification.

What distinguishes µODE is the composable ecological layer, and the colonization-resistance result shows why this is not a cosmetic difference. The mechanism that produces resistance (i.e. 7α-dehydroxylation of primary to secondary bile acids, the germination dose-response, and the antibiotic-driven collapse of the effector guild) is not encoded in the genome-scale reconstructions at all: the *C. scindens* GEM carries no bile-acid exchange reactions, because bile transformation is a rate-modifying ecological process rather than a balanced metabolic flux. Consequently, as configured with these GEMs alone (which carry no bile-acid reactions), engines that operate solely on the supplied models (i.e., COMETS, MICOM, SteadyCom, µbialSim) have no bile-acid axis and do not represent the phenotype. This reflects the content of the supplied models rather than an intrinsic limit of every engine: COMETS, for example, supports extracellular reactions with kinetic parameters and could represent the bile chemistry if it were manually encoded in the layout. µODE's contribution is to supply this chemistry as a composable ecology layer by default. Running the identical 5-member community under a steady-state community objective (MICOM cooperative trade-off) returns a single coexistence snapshot (all five members at $\mu \approx 0.32\ h^{-1}$) whose entire metabolite space contains zero bile-acid species (no time axis, no bile pool, and therefore no antibiotic-induced bile inversion). Conversely, plain dFBA without the ecology layer (i.e., the computation COMETS performs on the same community) collapses to a pathobiont bloom and yields no resistance (Table 3: Bray-Curtis 0.983). Only µODE, coupling the dFBA core to the composable bile/pH/pharmacokinetic layer, reproduces the calibrated physiological deoxycholate concentration (0.47 mM vs measured 0.59 mM), its uncalibrated antibiotic-induced collapse, and the resulting 44-fold shift in the simulated *C. difficile* susceptibility ratio. The comparison is deliberately even-handed: for pure steady-state composition MICOM outperforms µODE's greedy dFBA (121/122 coexistence; Figure 5 inset), which is precisely why the dynamic, ecology-aware regime is the complementary capability µODE adds rather than a claim of universal superiority.

### SUPPLEMENTARY FIGURES

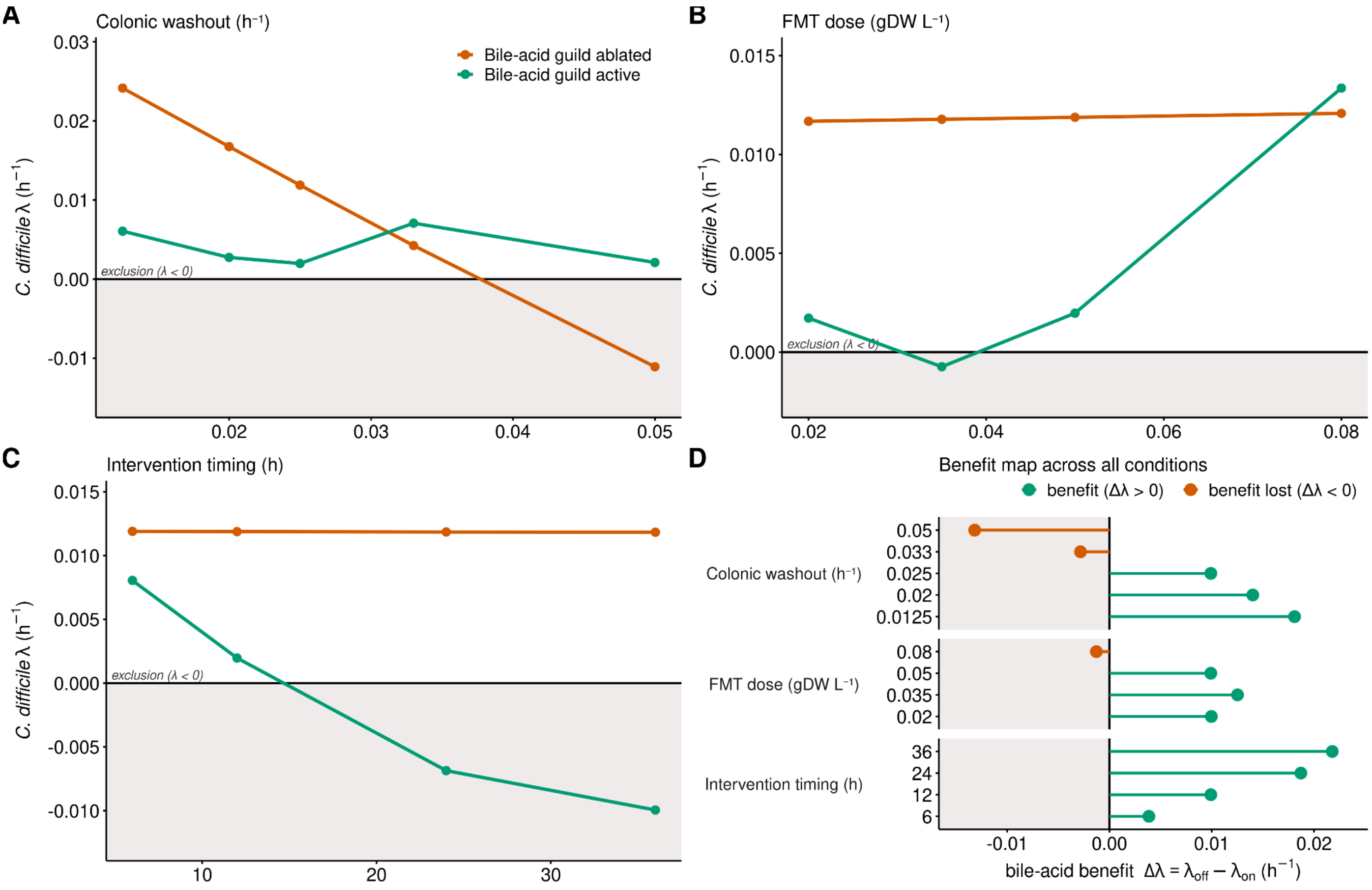


**Figure S1:** Sensitivity of the secondary-bile-acid arm of colonization resistance to three physiological parameters, evaluated on the defined five-member *C. difficile* community (colonic ModelSEED medium) at the same anchored operating point used in Figures 3-4 (community 7α-dehydroxylation rate vmax_bai=5, at which predicted deoxycholate matches the measured cecal concentration). For every parameter value the community was simulated twice, with the bile-acid ecology layer active and ablated, and the *C. difficile* endpoint rebound rate λ was recorded for each; the bile-acid benefit is defined as Δλ = λ(ablated) - λ(active), so positive values indicate that an intact bile-acid guild slows the pathogen. In all panels the shaded band marks pathogen exclusion (λ<0, i.e., net washout). Panel A – Colonic washout (dilution rate). With the guild ablated (vermillion), *C. difficile* rebounds falls monotonically as flow increases (λ from +0.024 $h^{-1}$ at +0.0125 $h^{-1}$ to -0.011 $h^{-1}$ at 0.05 $h^{-1}$), because at high washout bulk flow clears the pathogen irrespective of the bile-acid arm. With the guild active (green), rebound is held low throughout (λ ≈ +0.002 to +0.007 $h^{-1}$). The benefit, the vertical gap between the arms, is large across the physiological transit range (Δλ = -0.003 $h^{-1}$ at 0.033 $h^{-1}$, -0.013 $h^{-1}$ at 0.05 $h^{-1}$), where the two arms cross and washout, not bile, dominates. Panel B – FMT/inoculum dose (bolus biomass). The ablated arm is essentially insensitive to dose (λ ≈ +0.021 $h^{-1}$ across the range), whereas the active arm rises steeply at the largest bolus (λ from ≈ 0 at 0.02-0.05 gDW $L^{-1}$ to +0.013 $h^{-1}$ at 0.08 gDW $L^{-1}$). The benefit is preserved across the realistic dosing range (Δλ = +0.010, +0.013, +0.010 $h^{-1}$ at 0.02, 0.035, 0.05 gDW $L^{-1}$) and is marginally lost only at the extreme inoculum (Δλ = -0.001 $h^{-1}$ at 0.08 gDW $L^{-1}$), where sheer competitive biomass suppresses the pathogen independently of, and slightly in tension with, the bile-acid arm. Panel C – Intervention timing (transplant time). The ablated arm is flat (λ ≈ +0.012 $h^{-1}$ at all times: the pathogen blooms regardless of when an ineffective transplant lands), while the active arm falls with delay and reaches net exclusion for later transplants (λ = +0.008, +0.002, -0.007, -0.010 $h^{-1}$ at 6, 12, 24, 36 h). The benefit therefore holds across every tested timing and strengthens as the transplant is delayed (Δλ = +0.004, +0.010, +0.019, +0.022 $h^{-1}$), the interval giving the bile-acid guild more time to reconstitute the inhibitory pool before the pathogen escapes. Panel D – Benefit map. The bile-acid benefit Δλ for every condition, grouped by sweep; green marks conditions where the benefit holds (Δλ > 0) and vermillion where it is lost (Δλ < 0). Across all three axes the physiological operating point of Figures 3-4 (washout 0.025 $h^{-1}$, transplant at 12 h) sits well inside the benefit region. The benefit is thus robust throughout the physiologically realistic envelope and attenuates only at extreme washout or extreme inoculum, where bulk transport overtakes the rate-modifying ecology layer, a defined operating envelope rather than a universal effect.

### SUPPLEMENTARY TABLES

| Capability | µODE | COMETS | MICOM | SteadyCom | BacArena | µbialSim |
|---|---|---|---|---|---|---|
| **Temporal model** | Dynamic (dFBA, SOA) | Dynamic (dFBA) | Steady-state | Steady-state | Dynamic (agent-based) | Dynamic (dFBA) |
| **Community coupling** | Shared extracellular pool + shared proteome budget | Spatial grid + shared media | Community objective (cooperative trade-off, L2) | Balance community growth (equal µ) | Individual agents on a lattice | Shared pool (well-mixed) |
| **Spatial structure** | Optional reaction-diffusion | 2D/3D (mature) | – | – | 2D lattice (individual-based) | – |
| **Multi-kingdom** | Yes (taxon-agnostic solver) | No | No | No | No | No |
| **Bacteriophage predation** | Yes (Levin-Stewart ODE ↔ host biomass | No | No | No | No | No |
| **Composable non-metabolic ecology layers** | Yes (stackable, literature-grounded) | Manual (extracellular reactions; not composable layers) | No | No | Partial (diffusion only) | No |
| **Timed perturbation engine** | Yes | Limited (media refresh) | No | No | Partial | Limited |
| **Automated MAG→GEM→simulation** | Yes (CarveMe / gapseq backends) | No (manual GEMs + layout) | Partial (taxonomy→AGORA) | No | No | No |
| **Scales to hundreds of strains** | Yes (deterministic ODE) | Yes (heavier with space) | Yes | Moderate | No (agent tracking intractable) | Yes |
| **Solver / runtime** | Python; scipy/HiGHS LP oracle | Java + GLPK/Gurobi | Python/COBRA; OSQP/Gurobi/CPLEX | COBRA; CPLEX/Gurobi | R + sybil; GLPK/CPLEX | MATLAB (COBRA Toolbox/CellNetAnalyzer); GLPK/Gurobi |
| **Interoperability** | COMETS export | – | – | – | – | – |

**Table S1:** Capability comparison of community metabolic-modeling frameworks. Entries reflect the default, out-of-the-box capability of each framework at its cited version; several engines can support additional behaviors through manual configuration.

### MATHEMATICAL FORMULATIONS

This supplementary section details the explicit mathematical formalisms, algorithmic routing logic, and software architecture underlying the µODE framework. While the main text focuses on the emergent biological and ecological capabilities of the engine, the following sections provide the complete suite of ordinary and partial differential equations, constraint-based formulations, and validation metrics required for rigorous computational reproducibility.

#### Community dynamics

Uptake Kinetics and Stability Limits: In dFBA, the bound on each uptake reaction is a function of the prevailing extracellular concentration. The maximum rate at which member $i$ can take up metabolite $j$, at concentration $M_j$, is defined as:

$$v_{j,i}^{up} = min\left(f_{ij}\frac{V_{max,ij}M_j}{K_{m,ij}+M_j}, \frac{M_j}{X_i\Delta t}\right)$$

The first term represents saturable, concentration-dependent uptake, where $V_{max,ij}$ and $K_{m,ij}$ are the capacity and affinity, respectively, and $f_{ij} \in [0,1]$ is an optional ecological modifier (e.g., for oxygen gating). The second term provides the stability condition for the explicit scheme: it forbids any member from removing more of a metabolite than is present within a single time step $\Delta t$.

Community ODE Integration: Because members' intracellular fluxes are assumed to reach a quasi-steady state rapidly relative to environmental changes, their metabolic problems are solved independently within a step. Once every member has been solved, the community is advanced using a first-order explicit integration. Biomass $X_i$ and metabolite concentrations $M_j$ update according to:

$$\frac{dX_i}{dt} = \left(\mu_i - \delta - D\right)X_i, \qquad \frac{dM_j}{dt} = \sum_i v_{j,i}X_i + \phi_j - DM_j + r_j^{eco}$$

Where $\mu_i$ is the computed growth rate, $\delta$ is an optional first-order death rate, $D$ is a chemostat dilution rate, $\phi_j$ is the influx of nutrient $j$ from the medium, and $r_j^{eco}$ is any additional rate contributed by the ecology layer (e.g., microbial transformation of a bile acid). Biomass and concentrations are strictly held non-negative.

Emergent Cross-Feeding Inference: A directed interaction from a producer to a consumer, mediated by a shared metabolite, is reported if the producer's net secretion coincides with the consumer's uptake at any time $t$ during the simulation:

$$\exists t\colon v_{j,prod}(t) > \epsilon \wedge v_{j,cons}(t) < -\epsilon$$

The interaction strength is subsequently summarized by the maximal coincident secretion flux.

#### Kinetics and enzyme constraints

Extracellular Uptake Kinetics: In dynamic flux balance analysis (dFBA), the bound on each uptake reaction is dynamically coupled to the prevailing extracellular concentration. This saturable, concentration-dependent uptake is modeled using Michaelis-Menten kinetics:

$$v^{up} = \frac{V_{max} M}{K_m + M}$$

where $V_{max}$ is the maximum uptake capacity and $K_m$ is the half-saturation constant (affinity).

Parameter Inference and Tiered Assignment: To handle the scarcity of measured kinetic parameters in automated reconstructions, unmeasured parameters are assigned via a tiered approach:

- Knowledge-Based Defaults: Explicit priors using experimentally measured $K_m$ values for common substrates, with representative values applied elsewhere. Turnover numbers are set to the genome-wise median, characterizing a moderately efficient enzyme.
- Deep-Learning Refinement: For enhanced resolution, sequence-based and structure-based deep learning models predict turnover numbers and half-saturation constants, treated as priors with quantified uncertainty to refine reaction bounds.

Intracellular Enzyme and Proteome Constraints: Intracellular fluxes are constrained by both local enzyme availability and a global proteome budget. For each intracellular reaction $r$ with a known turnover number $k_{cat}$, the reaction rate $v_r$ is strictly bounded by:

$$|v_r| \le k_{cat,r}[E_r]$$

where $E_r$ is the effective abundance of the catalyzing enzyme.

To capture the competing demands for limited cellular resources, a global proteome budget is enforced across all enzyme-limited reactions:

$$\sum_r \frac{MW_r}{k_{cat,r}} |v_r| \le P$$

Each reaction's contribution is weighted by the mass of enzyme required to sustain it (proportional to its molecular weight $MW_r$ and inversely proportional to its turnover number $k_{cat,r}$). $P$ represents the fixed fraction of the total proteome allocated to these metabolic reactions. This constraint is integrated as a standard linear reformulation and is dynamically maintained throughout the simulation.

**Perturbations and interventions**

Perturbation Operators: Perturbations are applied to the metabolic constraints before the dynamic integration begins. For pathway attenuation, which models partial drug inhibition, the capacity of every reaction in a specified pathway is reduced by a defined efficacy parameter:

$$V_{max} \leftarrow (1 - efficacy)\, V_{max}\,, \qquad efficacy \in [0, 1]$$

Full efficacy (1.0) is mathematically equivalent to a targeted reaction knockout, which removes the specified reactions entirely by setting their bounds to zero. Member removal eliminates the organism's corresponding dimensions from the community state vectors entirely.

Discrete Event Scheduling for Interventions: Timed interventions are modeled as discrete state updates during the ODE integration. For an intervention scheduled at time $t_{int}$, the introduced organisms $i \in I_{donor}$ are initialized with $X_i(t) = 0$ for $t < t_{int}$.

At $t = t_{int}$, the community biomass vector is discontinuously updated using the defined inoculum bolus:

$$X_i\left(t_{int}\right) = X_i\left(t_{int}^-\right) + \Delta X_{i,inoculum}$$

For a transplant, the inoculum $\Delta X_{i,inoculum}$ is derived from the relative composition and total absolute size of the donor community. Donor organisms that are already present in the recipient community have their existing biomass reinforced by the inoculum rather than duplicated in the state space.

**Ecology layer**

pH and Short-Chain Fatty Acid Inhibition: Medium pH falls as acid accumulates and is modeled as a buffered titration. Each acid is partitioned into its charged and neutral forms via the Henderson-Hasselbalch relation, as only the undissociated form ($HA$) enters the cell:

$$pH = pH_0 - \frac{A_{total}}{\beta}, \qquad HA = \frac{A}{1+10^{pH-pK_a}}$$

Growth is scaled by an undissociated-acid inhibition term combined with a cardinal-pH window (ρ) that vanishes outside a viable range and peaks at the optimum:

$$g_i^{pH} = \frac{1}{1+\sum_a HA_a/K_{i,i}} \times \rho\left(pH;\ pH_{min}, pH_{opt}, pH_{max}\right)$$

Where $K_{i,i}$ denotes the sensitivity of member $i$ to the acid, allowing dense SCFA producers to suppress acid-sensitive pathogens.

Pharmacokinetics and Antimicrobial Killing: Drug concentration $C(t)$ is modeled via single-compartment pharmacokinetics with first-order elimination over a dosing schedule:

$$C(t) = \sum_{t_d \le t} dose \cdot e^{-k\left(t-t_d\right)}, \qquad k = \frac{ln2}{t_{1/2}}$$

This time-varying concentration drives a saturating ($E_{max}$) killing rate of the vegetative biomass ($\delta_i^{drug}$) of susceptible members:

$$\delta_i^{drug}(t) = E_{max}\frac{C(t)}{EC_{50}+C(t)}$$

Spores, residing in a separate biomass pool, are unaffected by this death rate.

Bile Acid Transformation Kinetics: Microbial transformation of host primary bile acids proceeds in two saturable steps, each first-order with respect to the biomass of the guild carrying the necessary enzyme (BSH or 7$\boldsymbol{\alpha}$-dehydroxylase):

$$taurocholate \rightarrow (bile\ -\ salt\ hydrolase) \rightarrow cholate \rightarrow (7\alpha\ -\ dehydroxylation) \rightarrow deoxycholate$$

**Reconstruction algorithms and quality control mechanics**

Kingdom-Aware Routing and Assembly Strategies: Reconstruction relies on kingdom-appropriate gene calling and modeling templates. For prokaryotes and archaea, genes are predicted directly from the nucleotide sequence. The default reconstruction is a fast top-down approach that carves a context-specific model from a curated universal template. An alternative bottom-up approach is available for specific taxa but operates in a separate namespace and at greater computational cost.

Downstream Structural Validation and Failure Isolation: Once reconstructed and gap-filled, every model is subjected to a battery of fast structural checks before being admitted to the community simulation:

- Biomass Viability: Verification that the model can successfully produce biomass on the defined medium.
- Mass and Charge Balance: Ensuring basic stoichiometric consistency.
- Energy-Generating Cycles: A strict test to identify artefactual cycles that would allow a model to generate ATP from nothing, which would otherwise invalidate dynamic predictions.

To support automated scaling across hundreds of genomes, the assembly step implements strict failure isolation. The results of the structural checks are aggregated into a per-run summary. Any model flagged as non-simulatable is silently excluded and logged for audit, ensuring that a handful of poor-quality MAGs do not abort a large-scale community simulation.

**Validation metrics and scaling policies**

Benchmark Scoring Metrics: A simulation is compared to user-defined expectations along three axes. An expectation need not be complete; unspecified aspects are simply not scored.

- Compositional Accuracy: Agreement between predicted and expected relative abundances, summarized by mean absolute error (MAE) and Spearman rank correlation (with expected-but-absent members counted as zero).
- Metabolite Accuracy: The root-mean-square deviation (RMSD) of predicted final concentrations from their expected values or ranges.
- Interaction Accuracy: Compares the inferred cross-feeding network to the expected one (at the producer-metabolite-consumer level) using precision, recall, and their harmonic mean (F1 score):

$$precision = \frac{|E_{sim} \cap E_{exp}|}{|E_{sim}|}, \qquad recall = \frac{|E_{sim} \cap E_{exp}|}{|E_{exp}|}, \qquad F1 = 2 \times \frac{precision \times recall}{precision + recall}$$

Abundance-Aware Reduction Policies: To compress large communities, µODE allows reduction under three explicit, user-defined policies:

- Top-N Retention: Retaining a fixed number of the most abundant members.
- Abundance Thresholding: Discarding members below a specific relative abundance threshold.
- Cumulative Fraction: Retaining the smallest set of members whose cumulative abundance reaches a chosen fraction of the whole community (e.g., capturing 99% of the initial biomass).

Post-reduction, the retained abundances are strictly renormalized. The choice of policy is a deliberate parameter left to the analyst, acknowledging the trade-off that rare members can occasionally drive essential specific interactions.
